# Starvation enhances bacterial survival through drying and rewetting at the single cell level

**DOI:** 10.64898/2026.01.12.697931

**Authors:** Jarek V. Kwiecinski, Georgia R. Squyres, Dani Or, Dianne K. Newman

**Affiliations:** Division of Geological and Planetary Sciences, California Institute of Technology, Pasadena, CA 91125; Division of Biology and Biological Engineering, California Institute of Technology, Pasadena, CA 91125; Department of Environmental Systems Science, Swiss Federal Institute of Technology, 8092 Zürich, Switzerland; Department of Civil and Environmental Engineering, University of Nevada, Reno, NV 89557

**Keywords:** osmolytes, desiccation, starvation, dormancy, soil

## Abstract

Soil bacteria are critical to agricultural productivity and play a central role in several biogeochemical cycles. These organisms frequently experience desiccation, which deprives them of access to both water and nutrients. Desiccation eliminates the aqueous connections between soil pores, removing pathways for nutrient diffusion. Well-studied yet resource intensive water stress responses like osmolyte synthesis may thus be impractical in unsaturated environments. Accordingly, we observed how the rhizobacterium *Pseudomonas synxantha* 2-79 responds to co-occurring water and nutrient limitation at the single-cell level to understand what role osmolyte synthesis may play in its desiccation response. We constructed a transcriptional reporter to track *P. synxantha*’s osmolyte response and collected extensive morphological and reporter expression data through experiments designed to mimic different rates and extents of soil drying. Only actively growing cells responded to an osmotic shock by synthesizing osmolytes: this response was not observed when we pre-starved bacteria. Soluble nutrient diffusivity in soil is restricted even when there is sufficient water to keep bacterial cells hydrated, so this pre-starved condition reflects gradual drying in which starvation precedes water stress. Despite the lack of osmolyte synthesis, prior starvation enhanced *P. synxantha*’s ability to recover from osmotic stress once water and nutrients were restored. These results suggest that cellular changes associated with the response to starvation that go beyond osmolyte synthesis play an important role in microbial desiccation tolerance.

**Significance Statement:** Soil bacteria play a vital role in agriculture by promoting plant productivity. Despite this, it is unclear how these organisms respond to desiccation, a common stress whose frequency is rising. Desiccation both dehydrates bacterial cells and eliminates the liquid water connections between soil pores that bacteria use to access nutrients. We studied how a model soil bacterium responds to simultaneous starvation and water stress at the single cell level, focusing on whether the bacteria synthesized osmolytes, small molecules that bacteria can accumulate to limit water loss. We found that starvation restricts osmolyte synthesis but enables bacteria to withstand more severe water stress. The starvation that accompanies desiccation may play a protective, rather than an antagonistic, role in microbial desiccation tolerance.

## Introduction

As Earth’s largest biome, drylands are home to ∼ 2 billion people and occupy ∼ 41% of the planet’s land area (1). Within these ecosystems, soil bacteria play a vital role in nutrient cycling, promote plant growth, and can confer enhanced drought tolerance to crops (2, 3). During dry periods, soil bacteria lose access to nutrients and experience decreases in soil matric water potential (Ψ), a quantity typically expressed with units of pressure (i.e. energy per volume) that captures the effects of capillarity and adhesion on the chemical potential of pore water (4). Upon rewetting, nutrient access is restored and cells become rehydrated; this drives a respiratory pulse known as the Birch effect (5).

Although studies of water stress often focus on its osmotic effects, in soils, fluctuations in the availability of water cause cells to experience both nutritional and biophysical stresses. While bacterial responses to water loss alone have been characterized (6), we know less about how bacteria adapt to the tandem water and nutrient challenges that occur during drying and rewetting in soil. Desiccation proceeds along a continuum (Figure 1). In a saturated soil, nutrients can freely diffuse through pore water and bacteria are in osmotic equilibrium with their environment. As soils begin to dry, nutrient diffusivity dramatically drops even though cells remain hydrated. Nutrient diffusivity is a critical variable for soil bacteria because they inhabit thin liquid films that, under even modestly dry conditions, are too small to support motility (7). Diffusion is thus the only means by which bacteria can access their substrates. As drying proceeds, liquid films become thinner and aqueous connections between pores disappear, increasing the diffusion path length between bacteria and nutrient sources (8). Consequently, effective solute diffusivity decreases significantly even when soils dry to relatively mild water potentials (9) that are easily tolerated by most bacteria (10). Eventually, the soil matric water potential decreases below the water potential of the cytoplasm, causing water to leave the cell (11). At more extreme water potentials that threaten cellular integrity and viability, nutrient transport essentially stops altogether (4, 9).

**Figure 1.**
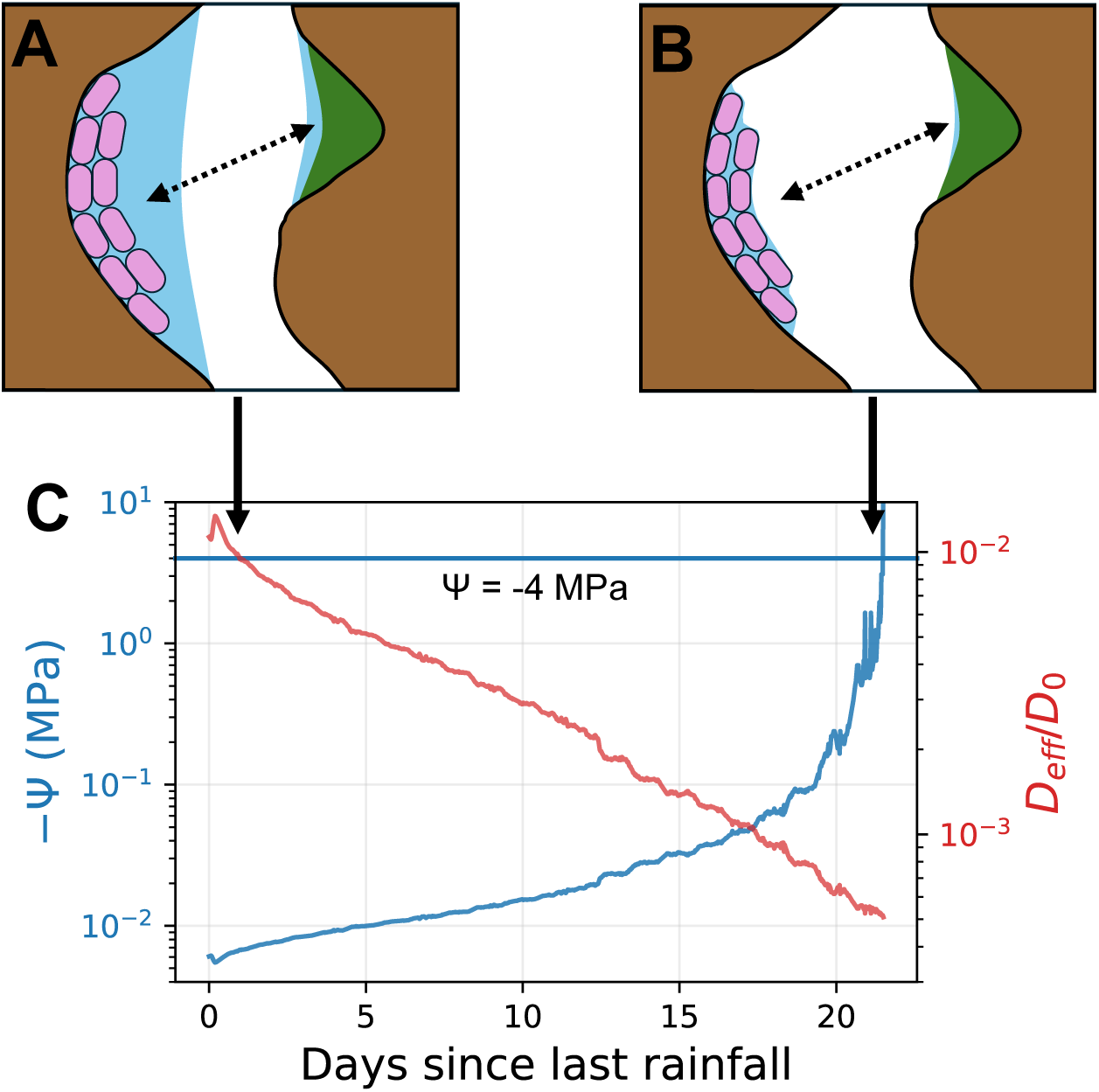
Solute diffusivity is dramatically reduced even in relatively wet soils. The cartoons show soil pores in different hydration states. In panel A, bacteria (purple rods) are well hydrated but lack a hydraulic connection to their substrate (green), meaning that they have no means to access nutrients from their surroundings. Panel B shows the same soil pore in a more advanced stage of drying at which bacteria begin to desiccate. Panel C shows how these dynamics play out in a real soil, supporting the notion that starvation precedes desiccation. Soil moisture data are from (70) and were taken from the San Joaquin Experimental Range NEON field site in the Sierra Nevada foothills following the last rainfall event of the 2023 rainy season. We computed the bulk averaged effective diffusivity of dissolved nutrients from this data according to the equation in (71). Even immediately after rainfall, the effective diffusivity was only ∼1% of the diffusivity in bulk water. This rain event did not saturate the soil, so there is limited hydraulic connectivity between any two points. Many soil microsites may look like the cartoon in A. The water potential at this time, as estimated from soil moisture using a van Genuchten retention curve (72), is less negative than any of the treatments we imposed in our experiments and would be easily tolerated by essentially all bacteria. The retention curve was parameterized using values for loamy sand, the texture at the field site (73), taken from (74). It was not until 3 weeks after rainfall that the water potential decreased to -Ψ = 4 MPa, the most stressful condition in our experiments (blue line in panel C). At this point, the diffusivity is more than an order of magnitude lower than where it began, and bacteria would have experienced considerable water stress as in panel B.

How do cells cope along this dynamic trajectory? In aqueous environments where nutrients are readily available, it is well established that bacteria adapt to water potential disequilibria by synthesizing osmolytes that allow them to retain cytoplasmic water while protecting proteins from damage (6, 12). Yet this strategy requires a significant resource investment (13) that may be impractical in the diffusion-limited soil matrix. This challenge is especially acute in the driest soils. For example, at a water potential -Ψ = 1.5 MPa, most plants can no longer extract water from soil, and the growth rate of most bacteria decreases (10). For bacteria to maintain turgor pressure under these conditions, osmolytes would have to account for roughly 6% of their biomass carbon, levels that are easily achieved by bacteria in liquid culture (14) but may be difficult to reach under nutrient limitation. In climates with lengthy dry periods, the water potential can decrease to as low as -Ψ = 30 MPa (15), and osmolytes would have to account for essentially all of microbial biomass for bacteria to avoid dehydration (13). The resource constraints and extreme levels of water deprivation that occur in very dry soil therefore seem to rule out osmolyte synthesis as a viable survival strategy. However, uncontrolled water loss damages the cell membrane, disrupts respiration, causes oxidative stress, and drives protein aggregation, severe consequences that threaten cell survival (16).

If osmolyte synthesis may not be used to protect bacteria over the entire desiccation trajectory, what other survival mechanisms might cells employ? Intriguingly, the cellular response to starvation itself may promote desiccation tolerance even if starvation constrains osmolyte synthesis. At the transcriptional level, nutrient depletion represses oxidative metabolism as well as nucleotide and rRNA biosynthesis, limiting oxidative stress and slowing overall transcription and translation rates (17). Reactive oxygen species accumulate under desiccation when the electron transport chain is disrupted due to membrane damage (16), so repression of oxidative metabolism under starvation may protect desiccated cells from oxidative damage. Starvation also upregulates genes that promote ribosome hibernation, inhibit DNA replication, and regulate carbon storage (18). These and other shifts in gene regulation drive broad changes to the overall structure of a bacterial cell under starvation. Cells become smaller and more spherical as they consume their cytoplasmic contents and undergo reductive division. At the same time, starvation makes the cell wall thicker (19), increasing its resistance to mechanical stress (20) and possibly limiting deformation to the cell envelope that occurs during dehydration (21). Consistent with these observations, prior starvation has been shown to help *Escherichia coli* better survive an osmotic shock (22), and nitrogen deprivation can somewhat enhance the survival of some cyanobacteria through desiccation (21, 23). The protective effects of nutrient deprivation may thus represent a key component of soil bacterial water stress survival that transcends osmolyte synthesis.

Accordingly, we set out to investigate how tandem nutrient and water limitation shape the bacterial water stress response. We chose to work with *Pseudomonas synxantha* 2-79, a model soil bacterium whose molecular and cellular responses to desiccation can be studied at the single cell level using live microscopy. We examined its response to (co)-variation in osmotic stress and nutrient deprivation to simulate different soil drying trajectories (Figure 1). We set out to test the hypothesis that the relative importance of osmolyte synthesis and starvation responses to desiccation tolerance is context dependent, with the optimal strategy determined by the rate and extent of drying cells experience.

## Results

We considered three scenarios to probe the effects of starvation and water stress in isolation and in tandem. First, we conducted experiments where we reduced the water potential in the presence of nutrients. Second, we simultaneously removed nutrients and lowered the water potential, simulating how bacteria become disconnected from their nutrient supply under desiccation. Finally, we starved bacteria before lowering the water potential, mimicking cases where nutrient diffusivity decreases before water availability restricts microbial activity (Figure 1). In all cases, we mimicked rewetting at the end of the experiment by restoring water and nutrients and tracked cell responses using a transcriptional reporter.

### Developing a transcriptional reporter to monitor osmolyte biosynthesis gene expression

To test the hypothesis that the osmolyte synthesis response to water stress is condition-dependent, we created a transcriptional reporter to monitor osmolyte biosynthesis gene expression. The first step in designing this reporter was to identify the osmolyte biosynthesis genes that *P. synxantha* upregulates the most under water stress. We therefore conducted an RNA-seq experiment in which we transferred a mid-exponential phase culture of wild type *P. synxantha* to a new medium with the same nutrient composition but augmented with 280 g/L polyethylene glycol (PEG 8000, average molecular weight = 8000 Da) to impose water stress. This medium was designed to mimic the average elemental composition of soil microbial biomass (24), and with PEG added, it had a water potential -Ψ = 2.66 MPa. PEG is used in studies of plant and bacterial water stress in place of salts because it is nonionic and therefore has no direct effect on membrane potential or ion homeostasis (25). Even though PEG of this size can be imported and metabolized by some pseudomonads (26), it has distinct effects on cellular ultrastructure and membrane composition from more permeating solutes like NaCl or PEG 200. These changes better match what happens to cells under desiccation (25). Following this treatment, we extracted and purified RNA before processing reads to evaluate differential gene expression between control and water stress treatments (Figure 2A).

**Figure 2.**
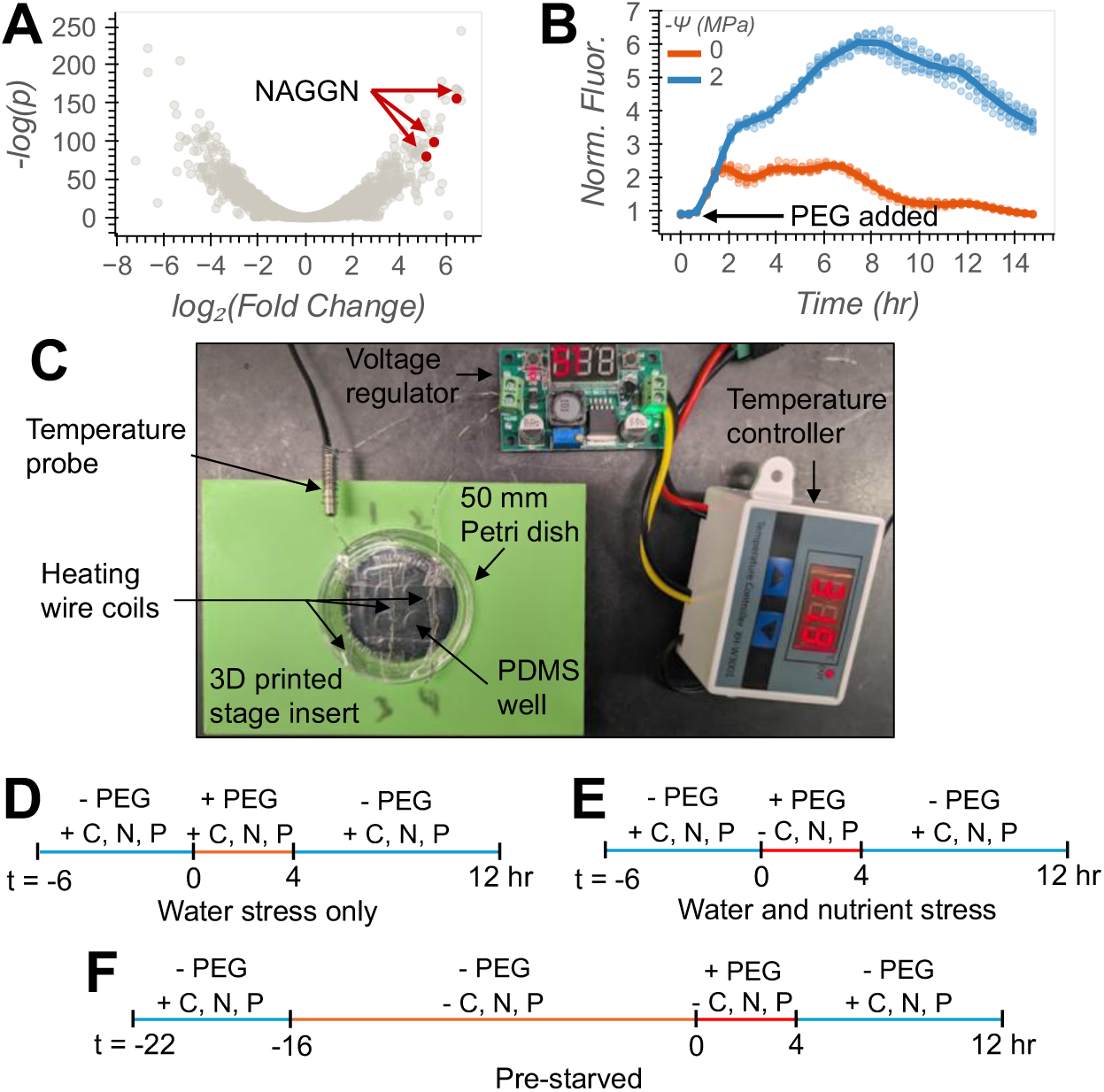
RNA-seq results, reporter verification, and setup for microscopy experiments. (A) Decreasing water potential with PEG alters *P. synxantha* gene expression. Genes with positive fold change values were upregulated upon exposure to PEG. Genes with smaller p-values (higher −log(*p*)) were covered by more sequencing reads or had larger fold changes, increasing confidence that these genes were in fact differentially expressed. NAGGN biosynthesis genes were the most upregulated genes for osmolyte synthesis and are highlighted in red. (B) The ratio of mNeonGreen to mApple fluorescence from our transcriptional reporter for osmolyte synthesis. mNeonGreen was expressed under control of a putative NAGGN promoter and mApple was expressed under control of the constitutive P_nptII_ promoter. Colors indicate the water potential the growth medium was adjusted to at *t* = 1 hr using PEG 1000. Points indicate the normalized fluorescence in individual replicates in a 96-well plate and lines show the mean. (C) Photograph of the experimental setup used to impose water and nutrient shifts over time and monitor cell behavior microscopically. Cells were grown in PDMS wells within a 50 mm glass-bottomed Petri dish affixed with a heated lid used to prevent fogging. (D-F) Timelines showing when PEG and nutrients were added and removed for each of our experimental conditions which included different combinations of water and nutrient stress.

The most up-regulated osmolyte synthesis genes were those associated with N-acetyl glutaminyl glutamide (NAGGN) biosynthesis (*ggnAB* and a putative peptidase, *P. synxantha* 2-79 locus tags C4K02_RS19705, RS19710, and RS19715, log_2_ fold changes of 5.2, 5.5, and 6.5). First discovered in *Sinorhizobium melitoti* (27), this dipeptide osmolyte is widely used by pseudomonads (28), especially the epiphytic plant pathogen *P. syringae* B728a (29). NAGGN production is known to be particularly resource intensive, and NAGGN synthesis can even harm some bacteria under osmotic stress because its production is so costly (29, 30). This made it an especially compelling target for our experiments, given our goal to investigate how starvation constrains the conditions under which osmolyte synthesis takes place.

After confirming that NAGGN production is highly induced during hyperosmotic stress, we designed a transcriptional reporter for its biosynthesis (Figure S1). First, we introduced the fluorescent protein mApple to a neutral chromosomal site downstream of a constitutive promoter (31). We transformed this strain with a plasmid harboring a putative promoter sequence for NAGGN biosynthesis (32) upstream of the gene for the fluorescent protein mNeonGreen. Attempts to put both the constitutive and NAGGN reporter constructs on the plasmid or both in the chromosome were unsuccessful, so we proceeded with this configuration. Having both constructs in the same strain allowed us to compute the ratio between mNeonGreen to mApple fluorescence (hereafter normalized fluorescence), a metric that accounts for factors beyond changes in gene expression that would affect reporter fluorescence. These factors include treatment effects on overall transcription and translation ability, fluorescent protein brightness, or changes in growth rate that would alter the rate at which fluorescent proteins are diluted into more biomass. Finally, we verified that this completed reporter construct was responsive to water stress by exposing exponential phase reporter cells to a water potential -Ψ = 2 MPa using PEG 1000 in a 96-well plate. Within 2 hours of exposure, the normalized fluorescence exceeded that of reporter cells that did not experience water stress (-Ψ = 0 MPa, no PEG added) (Figure 2B). PEG 1000 was used for this and future assays because of difficulties manipulating viscous PEG 8000 solutions. Though it is likely more membrane permeable than PEG 8000, PEG 1000 is still a larger molecule than solutes like sucrose that are commonly used for osmotic stress experiments. It is also uncharged, again avoiding confounding effects on ion homeostasis that are not relevant to the unsaturated environments represented by our experiments.

After verifying our reporter’s responsiveness to water stress via PEG addition, we monitored cell size, growth, and reporter activity over different conditions. We conducted these experiments using an apparatus adapted from (33) consisting of a glass-bottomed Petri dish with four PDMS wells (Figure 2C). The bottom of the wells was coated with an adhesive that held cells in place even when removing and replacing the fluid in the wells. This allowed us to monitor cell behavior through a series of nutrient and water potential changes, outlined in Figures 2D-F. In all cases, exponential phase reporter cells were first adhered to the bottom of the wells and allowed to adjust to the surface for six hours. Thereafter, we switched culture conditions by removing and replacing the full volume of the medium in the wells, changing the nutrient and PEG content as indicated. During our experiments, we collected single-cell phase-contrast and fluorescence microscopy images, segmented them to locate the boundaries of cells, and extracted single-cell morphological data, growth rates, and fluorescence intensities. Experiments were done in biological duplicate, so we studied how two different populations of cells responded to the same treatments. In each replicate, there were roughly 45-470 segmentation masks containing single cells or groups of cells present throughout the PEG treatments, with numbers varying between conditions. See Methods and Figures S5-S6 for details.

### Response of *P. synxantha* to reduced water potential in the presence of nutrients

We first observed *P. synxantha*’s cellular response to a decrease in water potential in the presence of nutrients. This condition represents the most well studied case of bacterial water stress in which changes in medium water potential are decoupled from nutrient availability. It thus served as a point of comparison to understand how nutrient deprivation drives deviations from the bacterial response to water stress alone. We tested water potentials -Ψ = 0, 1, 2, and 4 MPa, specified by PEG addition to the targeted water potential values, representing a realistic range of water potentials in desiccating soils (Figure 1C).

When the water potential was lowered to -Ψ = 1 MPa, cells continued growing, mounted a robust osmolyte synthesis response, and shrank very little despite the imposed hyperosmotic shock (Figure S2A). However, cells immediately reduced their growth rate to roughly two-thirds the value observed before PEG was added (Figure 3A). NAGGN biosynthesis genes were upregulated almost immediately, and both the normalized (Figure 3B) and mNeonGreen fluorescence (Figure S2C) reached an initial peak after 1 hour. The normalized signal began decreasing before PEG was removed, indicating that the rate of NAGGN biosynthesis gene expression had begun to decrease relative to the rate of constitutive mApple biosynthesis. The normalized reporter signal decreased more rapidly once PEG was removed. This is due to a combination of decreased NAGGN gene expression, as expected with the removal of water stress, and recovering cell growth rates (see increased growth rate after *t* = 4 hr in Figure 3A), increasing the rate at which the remaining mNeonGreen was diluted into more biomass.

**Figure 3.**
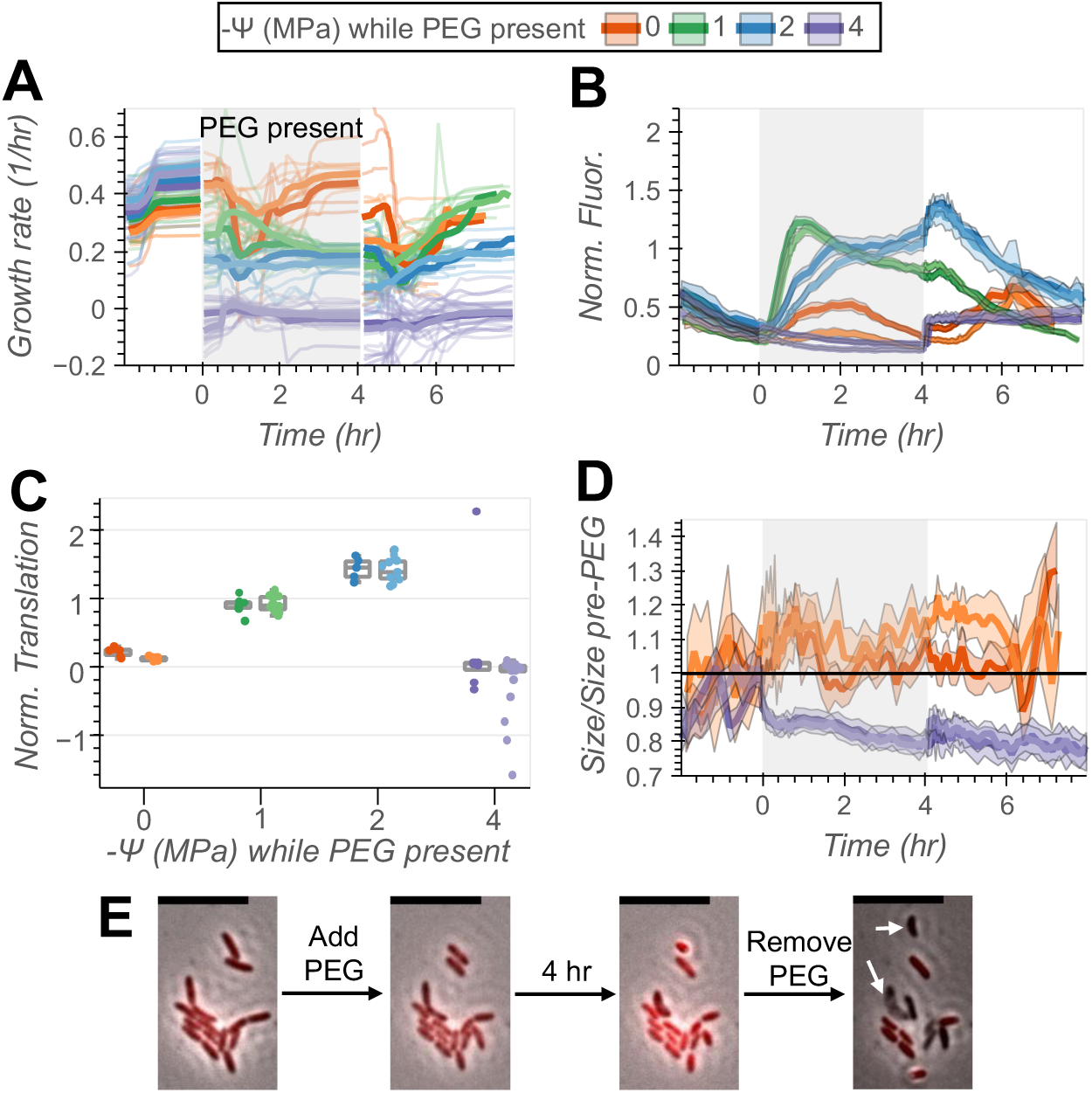
Cellular responses to water stress vary with the severity of the stress in *P. synxantha*. (A) Growth rate before, during, and after PEG exposure. PEG was present for the time range shaded in grey. Bold lines show the mean growth rate across all microcolonies tracked to measure growth (see Methods). Faint lines show growth rates for individual microcolonies. The number of microcolonies measured at each time point is displayed in Figure S7. Colors indicate different water potentials during PEG exposure, with darker and lighter shades distinguishing between replicate experiments. (B) NAGGN reporter fluorescence normalized by constitutive fluorescence. Lines represent the mean normalized fluorescence across all segmentation masks and ranges show 95% confidence intervals. The number of segmentation masks used to compute the mean fluorescence can be found in Figure S5. During PEG exposure, there was an average of 46-204 masks present, varying between water potentials and replicate experiments. (C) The ratio of total estimated mNeonGreen to mApple translation calculated using a model that accounts for photobleaching and growth dilution effects (see Methods). Points represent microcolonies. Higher values of this ratio correspond to greater overall NAGGN gene expression. Some values are negative for the -Ψ = 4 MPa cells because values of both integrated mNeonGreen and mApple translation were near zero (see Figure 6) and the model does not account for fluorescent protein leakage or degradation. (D) Size of reporter cells in the -Ψ = 0 MPa and 4 MPa treatments, normalized to their mean size in the 30 minutes leading up to PEG addition. Mean sizes displayed as lines were computed for segmentation masks filtered to contain only one cell (see Methods). The number of filtered masks used to compute mean sizes can be found in Figure S6. During PEG exposure, there was an average of 33-165 cells present, varying between water potentials and replicate experiments. (E) Images of one microcolony of reporter cells immediately before PEG addition, immediately after the water potential was lowered to -Ψ = 4 MPa, after 4 hours of exposure to this water potential, and immediately after PEG was removed. The phase contrast and control fluorescence layers are overlaid to show changes in cell morphology and membrane permeabilization leading to loss of mApple upon PEG removal. White arrows in the last image point to cells that lysed or were permeabilized. Scale bars are 10 µm. Red fluorescence data are all displayed on the same intensity scale.

*P. synxantha* also tolerated a shift to -Ψ = 2 MPa with only modest changes to cell morphology and growth. Though this treatment caused a greater initial decline in growth rate, cells from both the -Ψ = 1 and 2 MPa treatments converged on a similar growth rate equal to about half that observed in the untreated cells by the end of PEG exposure (Figure 3A). Cells shrank only slightly and regained their original size within 1 hour of PEG addition (Figure S2B), by which time NAGGN biosynthesis gene expression had already begun (Figures 3B and S2C). Reporter fluorescence began to increase later than in the -Ψ = 1 MPa treatment, suggesting a lag between the onset of water stress and osmolyte gene expression. Both the normalized (Figure 3B) and mNeonGreen fluorescence (Figure S2C) continued increasing until PEG was removed, so even though NAGGN biosynthesis gene expression began later, it continued for longer. We estimated total fluorescent protein production during the PEG treatment, applying a model that accounted for protein maturation kinetics, photobleaching, and dilution via growth (see Methods). The ratio of estimated mNeonGreen to mApple translation was highest in the -Ψ = 2 MPa treatment, so when accounting for differences in overall translation capacity, this water potential prompted the most osmolyte gene expression (Figure 3C). When nutrients are available, the milder -Ψ = 1 and 2 MPa conditions are clearly within *P. synxantha*’s ability to osmoregulate, and NAGGN biosynthesis represents a part of *P. synxantha*’s response to a modest shift in water availability.

Cells exposed to a water potential of -Ψ = 4 MPa followed a different trajectory. PEG addition immediately halted their growth, and growth did not resume even after PEG was removed (Figure 3A). The cells also shrank substantially, losing up to 15% of their original size within 1 hour of PEG addition (Figure 3D). Shrinkage continued at a slower rate for the rest of the 4-hour PEG treatment, and the cells ended at ∼ 80% of their original size before PEG was removed. Cells that survived PEG removal briefly increased in size before they shrank once again (see small increase at *t* = 4 hr in Figure 3D), and no detectable NAGGN gene expression occurred at any point in the experiment (Figures 3B and S2C). This is despite the fact that translation was still possible, as indicated by the increase in control fluorescence (Figure S2D) reflecting ongoing mApple translation during PEG exposure (purple points in Figure 6A, water stress only experiment). However, whatever translation occurs under these conditions is insufficient to recover cytoplasmic water during PEG exposure or to prevent widespread cell lysis upon PEG removal. Lysed cells were visible in images taken immediately after PEG removal (Figure 3E). Other cells were permeabilized and lost fluorescence, consistent with fluorescent proteins leaking into their surroundings. This was evident in the control fluorescence data, with sharp drops in fluorescence occurring immediately after PEG was removed (Figure S2D). Exponentially growing, un-adapted cells are thus not equipped to survive rapid, large shifts in water potential.

### Response of *P. synxantha* to simultaneous removal of nutrients and water

After establishing how *P. synxantha* responds to water stress in the presence of ample nutrients, we considered the opposite extreme of rapid drying in which cells lose access to nutrients and begin to dehydrate simultaneously. We observed *P. synxantha*’s cellular responses to a decrease in water potential and the simultaneous removal of nitrogen, phosphorus, and all carbon sources except for PEG. This condition provides no window for adaptation prior to the onset of tandem water and nutrient stress and therefore represents a worst-case scenario for soil bacteria. We again performed duplicate experiments, with one difference: the inocula for each experiment began at different growth rates prior to the onset of water and nutrient stress (see growth rates where *t* < 0 in Figure S3A and Figure S3B). This fortuitous difference revealed nuanced and interesting responses.

First, we examined cellular responses to nutrient removal in the absence of water stress (-Ψ = 0 MPa). In both inocula, growth rates decreased to similar levels upon nutrient removal, as expected under nutrient deprivation (red points in Figure 4C, right). In the faster-growing inoculum, cells were larger when entering the starvation treatment (Figure S3C) and shrank in size after 2 hours of nutrient deprivation (dark red line Figure 4A, top), consistent with the expected correlation between bacterial cell size and growth rate (34). Eventually, these cells were only slightly larger than the cells from the slower growing inoculum, which entered the starvation treatment at a smaller size (faint red line in Figure 4A, top). Cells under nutrient starvation made less mApple than when nutrients were never removed, consistent with reduced protein synthesis under nutrient limitation (0 MPa data points in Figure 6A), with similar mApple translation between inocula.

**Figure 4.**
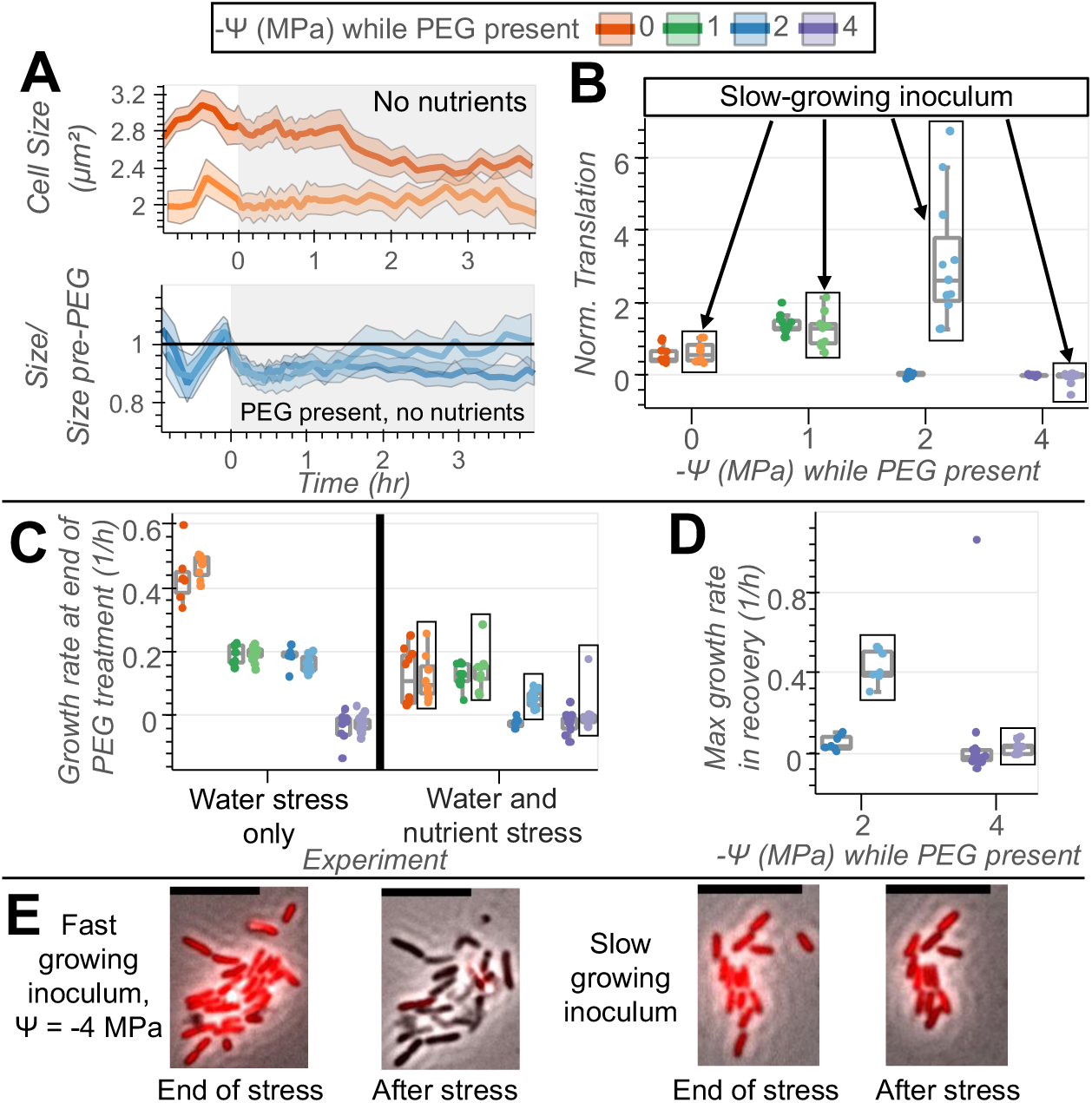
Cellular responses to the simultaneous removal of water and nutrients vary with growth rate prior to stress onset. (A, top) Mean size of reporter cells that experienced only nutrient stress beginning at *t* = 0 hr. Ranges show 95% confidence intervals. The lighter colored line shows means from the replicate experiment with a slower growing inoculum. During PEG exposure (grey shading), there was an average of 84 cells present for the replicate with a faster growing inoculum (replicate 1, dark colored line) and 33 cells present for the replicate with a slower growing inoculum (replicate 2). (A, bottom). Mean normalized size of reporter cells for which nutrients were removed and the water potential was reduced to -Ψ = 2 MPa at *t* = 0 hr. Cell sizes are normalized to the mean value in the 30 minutes leading up to PEG addition. During PEG exposure, there was an average of 97 cells present for replicate 1 and 43 cells present for replicate 2. Complete cell counts for the entire time series are in Figure S6. (B) The ratio of total estimated mNeonGreen to mApple translation calculated using a model that accounts for photobleaching and the dilution of fluorescent proteins by growth (see Methods). Points represent microcolonies, and points in boxes represent microcolonies that originated from the slower-growing inoculum. Higher values of this ratio correspond to greater overall NAGGN gene expression. (C) Mean growth rate in the last hour of PEG exposure, comparing the experiments with water stress only (left) and experiments where water and nutrients were simultaneously removed (right). (D) The maximum growth rate sustained for at least 0.5 hr for microcolonies in the -Ψ = 2 and 4 MPa treatments after PEG was removed and nutrients were restored. (E) Images of reporter cells at two different time points: 1) after 4 hours of simultaneous PEG exposure (-Ψ = MPa) and nutrient limitation and 2) immediately after PEG was removed and nutrients were restored for recovery. Cells on the left originated from the fast growing inoculum and cells on the right originated from the slow growing inoculum. The red overlay shows constitutive mApple fluorescence. The same intensity scale was used for all images and scale bars are 10 µm.

There were also only modest differences between inocula in the -Ψ = 1 MPa treatment. Just as when nutrients were maintained throughout water stress, the cells did not noticeably shrink immediately after PEG was added despite the hyperosmotic shock (Figure S3D). Growth rate decreased when PEG was added and nutrients were removed, but this growth rate matched that under nutrient removal only (compare green and red points in Figure 4C, right). In other words, water stress did not further reduce growth rate in this condition, even though water stress sharply decreased the growth rate in the previous experiments when nutrients were not removed (Figure 4C, left). The cells expressed NAGGN biosynthesis genes at high levels; in both replicates normalized mNeonGreen translation matched levels seen when nutrients were never removed (compare Figures 4B and 3C). This response was again rapid in onset but transient, with the normalized fluorescence decreasing rapidly after reaching its peak in 1 hour (Figures S3E-F). Differences in prior growth conditions had only subtle effects on *P. synxantha’*s ability to withstand nutrient stress occurring in isolation or in combination with mild water stress.

However, more extreme water stress amplified the differences between the inocula. The faster growing cells shrank by about 10% immediately after the water potential was lowered to -Ψ = 2 MPa, a similar extent to what we observed when nutrients were present (dark blue line in Figure 4A, bottom). These cells could not recover their original size, stopped growing completely (dark blue points in Figure 4C, right), and did not resume growth even after PEG was removed and nutrients were restored (dark blue points in Figure 4D). Even though the estimated amount of mApple synthesis was similar for both inocula and matched the levels seen with no PEG (blue points in Figure 6A, water and nutrient stress experiment), there was no detectable NAGGN gene expression by cells from the faster growing inoculum (dark blue points in Figure 4B and Figure S3E). Water stress was better tolerated by the slower growing cells, which also initially shrank by ∼10% but regained their original size within two hours of PEG exposure (light blue line in Figure 4A). Growth continued, albeit at a very slow rate (light blue points in Figure 4C, right), and the growth rate dramatically increased once PEG was removed and nutrients were restored (Figure 4D). NAGGN gene expression followed similar patterns to what we observed with nutrients present: gene expression at -Ψ = 2 MPa was delayed in onset but persisted for longer when compared to the response at -Ψ = 1 MPa (Figure S3F). The ratio of integrated mNeonGreen to mApple translation was also higher than in any other condition we tested (Figure 4B). Slower growing cells were better able to withstand and adapt to PEG addition.

We observed similar results for the -Ψ = 4 MPa treatment. This water potential was still difficult for *P. synxantha* to manage. First, in both replicates, cells shrank to a similar extent as when nutrients were present during water stress (Figure S3G). Subsequently, they could not grow while PEG was present (Figure 4C) or resume growth when water and nutrient stress ended (Figure 4D) and failed to express NAGGN biosynthesis genes (Figure 4B). However, removing PEG and restoring nutrients caused more cell lysis for the faster growing inoculum as indicated by a much larger decline in control fluorescence observed at the time of this transition (Figure S3H, see images in 4E). Again, the slower growing inoculum was better able to withstand large and rapid shifts in water potential.

### Response of pre-starved *P. synxantha* cells to water stress

Motivated by our observation that slower growing cells appeared to be advantaged under conditions relevant to extreme drying, we sought to explore the connection between prior nutrient limitation and water stress tolerance by pre-starving *P. synxantha* before adding PEG. This experiment approximates the starvation that is expected to precede dehydration in more gradually drying soils (Figure 1). Specifically, we deprived *P. synxantha* of nutrients for 16 hours prior to PEG addition, predicting that pre-starvation would enhance *P. synxantha*’s tolerance to large shifts in water potential.

In the short term, lowering the water potential had a similar effect on the pre-starved cells as it did in the other nutrient conditions. Cells initially shrank by a similar amount as when nutrients were present (compare Figures 5A and 3D). In our previous experiments, cells could not permanently recover any of their lost volume when shifted to a water potential of -Ψ = 4 MPa. However, with prior starvation, cells shifted to this water potential rebounded slightly in size within one hour of PEG addition (purple lines in Figure 5A). It is unclear how this recovery occurred, as there was no detectable NAGGN gene expression following PEG addition (Figures 5B and 6B). New gene expression may have simply been impossible during this period, as overall mApple translation was negligible (Figure 6A) and the control fluorescence even decreased while PEG was present (Figure 5C). Prior to PEG addition, the mNeonGreen fluorescence did increase, peaking after cells had been in the starvation medium for roughly 9-12 hours (*t* = −7 to −4 hr in Figure 5D). It was difficult to estimate total translation from fluorescence data during this period because our method requires growth rate data to account for fluorescent protein dilution. The growth rate, which by visual inspection appeared to be decreasing, could not reliably be estimated because cells began stacking on top of each other and later detached from the Petri dish surface in large numbers as starvation became severe (Figure S4A). However, the mNeonGreen fluorescence peak was highly variable between water potentials and biological replicates and coincided almost exactly with a peak in mApple fluorescence (Figures 5C-D). Therefore, the normalized fluorescence changed very little (Figure 5B), implying that the observed increase in mNeonGreen fluorescence was not driven by upregulation of NAGGN biosynthesis genes. Instead, changes that would affect both the mApple and mNeonGreen fluorescence in tandem likely explain this peak. Decreases in the growth rate during starvation would decrease the rate at which fluorescent proteins are diluted, and fluorescence can appear brighter because of vertical cell stacking. Consequently, we concluded that the extent of NAGGN biosynthesis by the starving cells was insignificant, so this osmolyte could not have contributed to *P. synxantha*’s survival through subsequent water stress and re-wetting.

**Figure 5.**
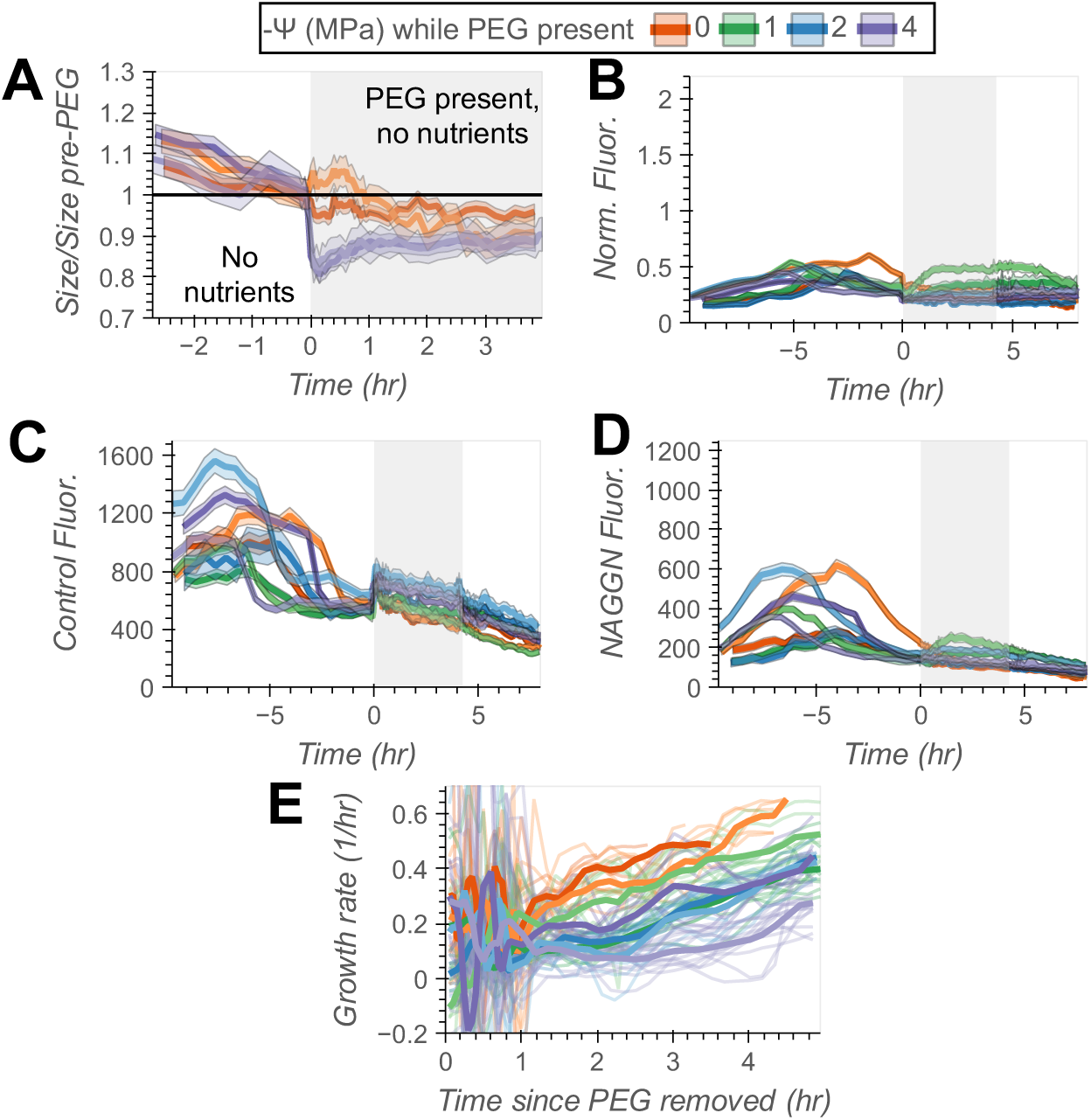
Pre-starved cells can recover from severe water stress without expression of NAGGN biosynthesis genes. (A) The normalized size of reporter cells after 16 hours of pre-starvation and over the course of 4 hours of simultaneous starvation and PEG exposure (grey shading). Lines show the mean cell size at each time point normalized to the mean size in the 30 minutes leading up to PEG addition. Colors show different water potentials, and the darker and lighter shades distinguish between replicate experiments. Ranges show 95% confidence intervals. During PEG exposure, there was an average of 145-355 cells present for the water potentials shown, with numbers varying between water potentials and replicates. Complete cell counts for the entire time series are in Figure S6. (B) The ratio of NAGGN reporter to constitutive fluorescence throughout the last 10 hours of pre-starvation, 4 hours of PEG exposure in the absence of nutrients, and 4 hours of recovery with water and nutrients restored. Lines show mean normalized fluorescence across all segmentation masks, including those with multiple cells. Ranges show 95% confidence intervals. During PEG exposure, there was an average of 128-469 masks present, varying between water potentials and replicate experiments. Complete mask counts for the entire time series are in Figure S5. (C) Constitutive mApple fluorescence during the same period, with the same number of masks used to compute averages. (D) mNeonGreen fluorescence, which peaked at the same time as mApple fluorescence during pre-starvation. (E) Mean microcolony growth rates after PEG was removed and nutrients were restored. Bold lines show the mean growth rate across all microcolonies, and faint lines show growth rates for individual microcolonies. The number of microcolonies measured at all time points is displayed in Figure S7.

Even though the pre-starved cells could not mount a transcriptional response to any decrease in water potential, they were better able to survive the transition back to unstressed conditions. The -Ψ = 4 MPa cells, which lysed after removing PEG in the other treatments, remained intact (Figure S4B) and showed no acute loss of fluorescence once PEG was removed and nutrients were restored (Figure 5C). These cells were later able to resume growth, a testament to their enhanced resilience (Figure 5E). This recovery was heterogeneous, as some cells began to elongate and divide while others recovered little in size (see increased variance in cell size distributions in Figure S4C and images in S4B). Still, the overall trends in growth and lysis demonstrate that prior stress exposure hardened *P. synxantha* cells against fluctuations in water potential, suggesting a role for starvation adaptation in bacterial survival through soil drying and rewetting cycles.

## Discussion

In this study, we investigated how a model soil bacterium responds to desiccation by (co)-varying the level of water stress and nutrient deprivation it experienced. This combination of stressors reflects the reality in soil environments that desiccation both deprives cells of water and restricts nutrient transport. We focused on *P. synxantha*’s ability to synthesize the osmolyte NAGGN under these conditions, as osmolyte synthesis is a well-known but resource intensive response to water stress. We studied this response at the single cell level in order to gain detailed insights into the effects of water and nutrient stress on both *P. synxantha’*s morphology and physiology. We hypothesized that NAGGN synthesis might be impractical under conditions simulating resource-limited dry soils. Our data indicate that while growing cells can make NAGGN while contending with water and nutrient stress, growth-arrested cells do not invest in NAGGN production. Indeed, cellular changes beyond osmolyte synthesis that accompany the response to starvation, like changes in envelope rigidity, may thus play an even more important role in soil bacterial desiccation tolerance.

If NAGGN synthesis is generally restricted to growing cells, as we observed for *P. synxantha,* the utility of this strategy may be limited for many soil bacteria. Isotopic measurements of microbial growth show that most taxa stop growing as soil dries out. Growth continues only for a small subpopulation consisting largely of Actinobacteriota that can continue growing at a similar rate as they did under moist soil conditions (35). These data suggest that dormancy is an option for many bacteria faced with desiccation, and strategies that enable bacteria to remain active through drying may be unnecessary or even unpreferred. Any active response to drying may instead serve to slow the onset of desiccation and preserve aqueous pathways for nutrient transport, both functions that can be achieved by producing extracellular polymeric substances (36, 37). Further experiments are needed to determine if osmolyte synthesis is essential for members of the growing subpopulation to remain active under dry conditions.

Moreover, our findings suggest that NAGGN synthesis may not adequately prepare soil bacteria for re-wetting, which unlike drying, occurs immediately in the natural environment. Rewetting is what ultimately killed *P. syxnantha* in our -Ψ = 4 MPa experiments when cells were not adequately adapted, and the cells that did survive did not express NAGGN biosynthesis genes in response to PEG addition. Stable isotope probing data show that the Actinobacteriota that continue growing in dry conditions (35) decrease in abundance following re-wetting and do not resume growth for 24 hours (38). Instead, the Pseudomonadota, including the Gammaproteobacteria of which *P. synxantha* is a member, dominate early growth. Cells that remain dormant during dry periods may be better prepared for rewetting, supporting their rapid recovery despite the potential costs of resuming growth from a dormant state. These dynamics have implications for soil carbon cycling because a significant pulse of microbial respiration follows soil re-wetting, a phenomenon known as the Birch effect (5) that is driven by the restoration of water and diffusive nutrient transport (39). These respiration pulses can account for up to 60% of net CO_2_ released from some dryland ecosystems during their dry season (40). Dormancy under dry conditions may enable greater participation of certain members of the microbial community in post re-wetting respiration, and the taxa that remain active during dry periods may contribute osmolytes and necromass that help fuel it.

Osmolytes are often detected in microbial biomass extracted from dry soil (41, 42). However, these compounds are found at bulk concentrations too low to balance the low water potential of the soil and prevent cytoplasmic water loss (42). Population averaged data, such as that collected in these studies, could miss variations in osmolyte production between growing and non-growing organisms that experience varied nutrient and water conditions at the microscale. Perhaps osmolyte synthesis is unevenly distributed within soil microbial populations. Alternatively, NAGGN is not the only osmolyte produced by *P. synxantha*, and other osmolytes that are better able to promote protein stability (43) may play a greater role not by limiting cytoplasmic water loss, but by protecting cellular machinery from desiccation damage (42). Trehalose is a common osmolyte that can prevent proteins from unfolding (43) and is strongly associated with long term desiccation survival in yeast (44). Unlike what we observed for NAGGN, it is also produced under starvation by some bacteria and yeast (45, 46). However, trehalose is not required for desiccation survival on the short time scales we addressed in our study, and trehalose synthesis alone cannot fully explain the desiccation tolerance of starved yeast cells (44). Further research is needed to characterize which osmolytes are relevant to desiccation tolerance, and to identify the specific effects of starvation that contribute to desiccation tolerance independent of osmolyte synthesis.

Like desiccation stress, starvation fundamentally remodels the bacterial cell. Respiration and translation are inhibited (17) while cells become smaller and the cell wall gets thicker (19). These structural and transcriptional changes do not appear to limit the loss of cytoplasmic water following a hyperosmotic shock, as the pre-starved cells in our experiments still shrank considerably immediately after PEG was added. However, these acute effects of water stress are clearly not predictive of *P. synxantha*’s overall resilience, and our findings stand in agreement with prior studies showing that starvation enhances recovery from salt stress (22) and even desiccation (23). Our study expands upon this work by characterizing the landscape of cellular responses to (co)occurring water and nutrient stresses, including changes to osmolyte synthesis, protein translation, cell size, and growth rate. Moreover, our work brings these single-cell responses at a high time resolution into the context of dynamic drying and rewetting soils.

In summary, our work demonstrates that the utility of NAGGN biosynthesis is conditional, and that a transition to low growth rate prior to the onset of water stress can also be protective. In soil, this transition is likely to occur through starvation, but the particular physicochemical properties that make starved cells more desiccation tolerant remain poorly understood. Increases in cell envelope strength under starvation may buffer *P. synxantha* cells against future shifts in water potential, allowing the organism to survive rewetting. In metabolically inactive cells, the cytoplasm can behave like a colloidal glass rather than a liquid (47), a phase change that also occurs under desiccation and slows the diffusion of biomolecules, protecting them from damaging cross-linking reactions associated with macromolecular crowding (48). Evidence for this phase change in starved cells is mixed (49), so further research is needed to identify any biophysical commonalities between desiccation and starvation that might explain starvation’s protective effects. The biophysical changes that accompany real desiccation, and not simply osmotic stress, have not been well explored in bacterial cells. Studying these changes may reveal traits beyond canonical water stress response pathways that confer desiccation tolerance by optimizing certain aspects of cellular ultrastructure. Identifying these traits would improve our understanding of dormant cell biology and is a priority for future research.

## Materials and Methods

### Strains and culture conditions

All strains were derived from *Pseudomonas synxantha* 2-79. This organism was isolated from a dryland wheat field in eastern Washington (50), where it defends wheat roots from fungal infection by producing antifungal phenazine compounds (51). Across the region, its relative abundance is highest in non-irrigated, drier sites (52). Unless otherwise indicated, cells were grown at 30⁰C with shaking in a defined medium (hereafter “SR”) intended to mimic the stoichiometry of soil microbial biomass (see composition in Table S1). The medium was also designed to have a low osmolarity and a pH that matched that of 1/2-21C, another low osmoticum medium used in prior studies of *Pseudomonas* osmotic stress response (c.f. (25, 29, 31)). For experimental treatments intended to impose nutrient starvation, we removed all sources of carbon (other than PEG), nitrogen, and phosphorous from SR and added more NaCl and MOPS buffer to match the osmolarity and ionic strength of SR, creating the medium SRM. Unless otherwise indicated, polyethylene glycol 1000 (PEG) was used to impose water stress at concentrations of 140, 209, and 300 g/L, corresponding to water potentials of -Ψ = 1.26, 2.00, and 3.65 MPa as measured by a WP4C dew point psychrometer. In the figures, these treatments are rounded to approximate water potentials of -Ψ = 1, 2, and 4 MPa respectively. Cells in the treatment labelled 0 MPa never received PEG. Tetracycline was used at a 15 µg mL^−1^ concentration (Tet15) when required for selection, or during all experiments with the reporter strain to maintain the reporter plasmid. *Escherichia coli* DH10*β* was used as a cloning host strain and gentamycin was used at 20 µg mL^−1^ (Gen20) with this strain for selection.

### RNA sequencing

We picked two colonies of wild-type *P. synxantha* from an LB agar plate and grew them overnight in separate 5 mL volumes of SR. Then, we diluted these cultures to OD 0.1 and grew them to OD 0.4-0.6, or mid-exponential phase. We split each culture into halves, one that was washed twice with SR and another that was washed with SR augmented with 280 g/L PEG 8000 (centrifugation done at 6800g for 5 minutes). After returning the cultures to 30⁰C with shaking for 30 minutes, we immediately pelleted the cells and performed RNA extractions as described in (53). We verified RNA integrity using an Agilent Bioanalyzer and performed rRNA depletion using a Ribo-Zero plus rRNA depletion kit (Illumina). Then, libraries were prepared using a NEBNext Ultra II RNA-seq kit (New England biolabs) and sequencing was done on an Illumina HiSeq2500 with a 2 × 50-bp configuration. Finally, we used Bowtie 2 (54) to align reads and featureCounts (55) to count gene expression. The R package DESeq2 (56) was used to compute the reported gene expression fold changes and p-values. Fold changes and computed p-values for all genes are available in Dataset S1, and raw reads are available at https://www.ncbi.nlm.nih.gov/bioproject/PRJNA1397269 (Dataset S2).

### Construction and validation of NAGGN biosynthesis reporter

After identifying NAGGN biosynthesis as a relevant target for further analysis, we designed a transcriptional reporter for the expression of the NAGGN biosynthesis genes. First, we used PCR and Gibson assembly to generate a plasmid containing the promoter sequence upstream of *ggnA* identified by (32), a synthetic ribosome binding site (RBS, (57)), and the gene for the fluorescent protein mNeonGreen (58). This plasmid was built on a modified pPROBETT backbone (59) adjusted to remove redundant terminators included in the original backbone sequence and containing an origin of replication that is functional in *Pseudomonas* (sequence in Dataset S3). We electroporated the plasmid directly into competent *P. synxantha* cells prepared according to (60). After 2 hours of recovery in Luria broth (LB) at 30⁰C, we plated transformants on LB Tet15 plates and allowed colonies to grow at 30⁰C overnight. Then, a subset of colonies was picked for sequence verification. Plasmids were purified using a New England Biolabs Miniprep kit from cultures derived from these colonies and sequenced by Plasmidsaurus. A plasmid with the correct sequence was stored at −20⁰C in ddH_2_O for future cloning.

Because osmotic stress alters the overall capacity of a cell to synthesize proteins (61), we also developed a constitutively fluorescent strain to account for this effect when considering changes in reporter fluorescence intensity. We chose the constitutive promoter for the neomycin phosphotransferase on the Tn5 transposon (P_nptII_), as this promoter was used as a positive control for gene expression in a prior study of *Pseudomonas* water stress response (31). We first attempted to add P_nptII_, another synthetic RBS, and the gene for the fluorescent protein mApple (62) to the plasmid harboring our NAGGN reporter construct. However, transforming this plasmid into *P. synxantha* significantly reduced its growth rate so we instead inserted a single copy of the constitutive reporter construct into a neutral site on *P. synxantha*’s chromosome. We used Gibson assembly to insert the construct into the pJM220 backbone (63) and transformed this plasmid into electrocompetent *E. coli* DH10*β* prepared using standard methods (64). After plasmid purification and sequence verification, we then transformed this plasmid and the helper plasmid pTNS1 (65) into electrocompetent *P. synxantha*. pTNS1 contains an integrase that catalyzed insertion of our constitutive reporter construct into the single *att*Tn7 site on *P. synxantha*’s chromosome as described in (65); pTNS1 and the pJM220 reporter plasmid are non-replicating in *P. synxantha* and so were not retained after outgrowth. A similar approach was used to generate a strain with mNeonGreen constitutively expressed under the control of the P_A1/04/03_ promoter (66) for photobleaching characterization. We also attempted to insert our NAGGN reporter construct into the *att*Tn7 site following the same strategy, but the level of expression driven by the NAGGN promoter was too weak to be detected when it was present only as a single copy. Thus, we transformed the constitutively mApple-producing red fluorescent strain with the NAGGN reporter plasmid described above, producing the “dual reporter” strain used for all subsequent experiments. Strains, plasmids, and primers for all cloning are listed in Table S2.

Finally, we conducted a plate reader experiment to verify that our reporter strain responds to water stress. We grew the strain overnight to stationary phase in SR at 30⁰C before diluting this culture to OD 0.1 and regrowing it to mid-exponential phase (OD 0.4-0.6). Then, we distributed this culture in 200 µL aliquots in a 96-well microtiter plate and recorded the OD_600_, red fluorescence (excitation = 561 nm, emission = 610 nm), and green fluorescence (excitation = 485 nm, emission = 535 nm) every 20 minutes for 1 hour using a Tecan Spark 10M plate reader to establish baseline activity. Incubations were done with shaking at 30⁰C. The first hour in Figure 2B shows data from this incubation used to establish a baseline level of reporter fluorescence. Separately, we prepared a plate with wells half filled with either SR or SR augmented with 400 g/L PEG 1000, roughly twice the concentration required to achieve a final water potential of -Ψ = 2 MPa. We transferred 100 µL of the cultures from the first plate into the plate with the PEG amended media, mixed thoroughly via pipetting, and placed this new plate into the plate reader to record the response to this hyperosmotic shock The lines in Figure 2B after PEG addition show data for the culture diluted 1:1 into SR (-Ψ = 0 MPa) or into SR + PEG 1000 (-Ψ = 2 MPa).

### Microscopy under water and nutrient stress

We began each water and nutrient stress experiment by preparing a mid-exponential phase culture of the reporter strain as in our reporter tests above. We also prepared a polydimethylsiloxane (PDMS) well plate using a method adapted from (33). Briefly, we prepared 20 × 20 mm squares of 2 mm thick PDMS and used a biopsy punch to create four 8 mm diameter wells in the PDMS. Then, we cleaned the PDMS and a 50 mm glass-bottomed Petri dish with 70% ethanol, 100% isopropanol, and 70% ethanol sequentially and plasma bonded the PDMS to the Petri dish. Immediately after plasma bonding, we added 50 µL of Corning CellTak adhesive solution prepared according to manufacturer instructions to each well. After letting sit for 20 minutes and rinsing out the CellTak solution, we diluted the mid-exponential phase culture to OD_600_ = 0.0015 in SR Tet15 and added a 0.5 µL droplet of culture to the center of each well. To limit evaporation, we also added 1 ml of SR with 15 µg L^−1^ tetracycline (SR Tet15) to the space surrounding the PDMS block inside the Petri dish. We glued the Petri dish to a custom 3-D printed stage insert and installed a lid with heating wires maintained at 32.5 - 33⁰C using a temperature controller and voltage regulator set at 1.5V. This temperature setting limited condensation on the lid of the Petri dish without preventing growth of the underlying cells. Cells were allowed to settle out of the small droplet for 10 minutes, during which time they became attached to the CellTak-coated glass at the bottom of each well. We began each experiment by filling the wells with 90-100µL of SR. At this point, imaging began. All microscopy experiments were done in duplicate.

Images were acquired using a Nikon Eclipse Ti2e microscope equipped with a 4.2 megapixel, 16-bit Orca Flash4.0v3 camera, a Lumencore Spectra X light source, and an Okolab H201 temperature control unit held at 30⁰C. We used a 40x 0.95 NA Ph2 objective with an additional 1.5x intermediate magnification tube lens activated for acquisition. In addition to phase contrast images, we collected mNeonGreen fluorescence images using 488 nm LED excitation with a 535/40 nm emission filter, and mApple fluorescence images using 561 nm LED excitation with a 620/60 nm emission filter. To ensure an adequate sample size of cells, we selected three fields of view for monitoring from each of the wells. FOVs were selected to contain a few to a dozen cells with adequate separation so that we could track the activity of microcolonies that developed from parent cells present at the beginning of the experiment. We imaged the cells every 10 minutes for six hours before beginning the next stage of the experiment. The purpose of this initial adjustment phase was to allow cells to adapt to the PDMS well plate and microscope environment before proceeding. From here, each experiment proceeded differently according to the type and sequence of imposed stresses; timelines of each experiment are presented in Figures 2D-F. In the paragraphs below, boldface text corresponds with the names of the experiments used in Figures 2-F and in figures 4C and 6A-B.

For the experiments with **water stress only,** we prepared SR medium augmented with the aforementioned PEG 1000 concentrations to impose water stress, along with a no PEG added control (-Ψ = 0 MPa condition). Then, well-by-well, we removed the existing SR medium, washed once with the target treatment medium, and refilled the well with this treatment medium to start the stress phase of the experiment. Because the Petri dish was glued to the stage insert, we could do this without moving the Petri dish to continue following the same FOVs selected above. Throughout this process and until an hour thereafter, we imaged every 3 minutes to capture any rapid osmotically driven changes in cell size that immediately followed PEG addition. We then reduced our imaging frequency to every 10 minutes and continued imaging for 3 more hours, meaning that cells were exposed to PEG-induced water stress for 4 hours in total. Finally, we resuscitated our cells by replacing all the media with SR. We again collected images every three minutes for one hour at the start of resuscitation before imaging every 10 minutes for 7 more hours.

For the experiments in which **water and nutrient stress** began at the same time, the experiment proceeded as described above, except that during the water stress, SRM starvation medium augmented with PEG was used.

Finally, for the **“pre-starved**” experiments where nutrients were absent both before and during water stress, we included an additional starvation phase after adjustment and prior to the PEG exposure. After the adjustment phase, we replaced all media with the SRM starvation medium to initiate nutrient stress. During this pre-starvation phase, we imaged every 30 minutes for 16 hours, as changes in cell size and reporter fluorescence were more gradual under this condition. We then replaced the media again with SRM augmented with PEG to initiate water stress, as above, and the rest of the water stress and recovery phases proceeded as outlined in the previous experiments.

### Segmentation and analysis of fluorescence images

To measure cell growth, cell size, and NAGGN biosynthesis reporter activity, we performed image segmentation. During these experiments, cells were initially seeded at low density, but over time they grew into small clusters, which we call microcolonies. We present analysis at the single cell, cell group, and microcolony levels. We limited our analysis to cells that were descendants of a parent cell that was present from the start of the experiment. This gave us information about the reporter’s response to all events in the experiment and limited our focus to cells that were always surface attached and therefore experienced similar conditions. We began by extracting tiff files for each frame from the nd2 files generated by the microscope and registered these images using the MATLAB package OmniSegger (67). Then, we manually selected one to five bounding boxes from each field of view containing a microcolony that developed over the course of the experiment. These boxes exceeded the final area of the microcolony so that its total area could be tracked over time.

After cropping the tiffs, we generated segmentation masks for cells within each microcolony using Omnipose, a deep neural network for image segmentation (68). We used the phase contrast images for this task with Omnipose’s pre-trained model “bact_phase_omni.” Segmenting individual cells was challenging because cell density within the microcolonies was high. We therefore limited our attempts at segmentation to time points when most cells in the microcolony were visibly distinguishable for each other. This is why some traces for reporter fluorescence and growth rates end earlier than others, especially in the milder stress conditions where microcolonies could grow larger and denser. Nonetheless, while some of the generated segmentation masks contained only one cell, many contained multiple. When analyzing our fluorescence data, we calculated the mean background subtracted fluorescence intensity for all the masks, giving us one fluorescence value per cell or small group of cells. We then plotted the mNeonGreen and mApple fluorescence, as well as the ratio between these two values, averaged across all the masks in a given PEG treatment. 95% confidence intervals for means were computed using the t distribution unless otherwise indicated. The number of masks used for these calculations at each time point is reported in Figure S5. Complete code for plotting and image analysis, as well as a list of required Python packages, are available at https://github.com/jkwiecinski0/NAGGNImageAnalysis (Software S1).

### Measuring cell size

To estimate cell size, we needed a method to filter for masks that contained only an individual cell. To do this, we separated each experiment into different stages where there were large visible differences in cell morphology. Though the exact definitions of these stages differed between experiments (see notebooks/filter.ipynb and src/mask_filter.py in Software S1), the -Ψ = 4 MPa cells were almost always analyzed separately from the other populations. Data from the adjustment period before PEG addition was also separated from later time points. Then, from each stage, we randomly sampled 100 masks and recorded the number of cells that fell within each mask. Next, we computed several morphological parameters including the length, width, and area occupied by the masks as well as the minimum curvature of the mask perimeters. We used principal component analysis to aggregate all of this information into two parameters and applied constrained optimization to find bounds on the principal components that effectively selected for masks containing only one cell. In our optimization approach, we sought to find bounds that maximized the fraction of the selected masks that contained only one cell. To ensure that the selected masks were still representative of the data, we required that our bounds included 95% of the masks from our random sample containing only one cell. Though these criteria could not always be met, the fraction of masks with one cell and the fraction of one-cell masks contained within our selected bounds were always greater than 84%. Values of the relevant principal components were then computed for the entire dataset and masks with more than one cell were filtered out using the optimized bounds. Figure S6 shows how many cells were left over at each time point after we performed this filtering. The means and confidence intervals for normalized size are derived from the filtered masks.

### Measuring microcolony growth rates

Segmentation quality was insufficient to track individual cells and compute single-cell growth rates. However, it was possible to track microcolonies of cells that grew from individuals at the start of each experiment and to measure their area over time. It was rare for cells to suddenly appear from elsewhere, so we did not account for the effect of these infrequent events to inflate the measured microcolony area. Conversely, several cells did not remain adhered to the bottom of the wells, so it was necessary to account for cell detachment by first detecting when cells left the field of view. We could detect these events as large, stepwise decreases in the area of the microcolony between adjacent frames and correct for them as needed. Specifically, we took the difference between images containing segmentation masks at adjacent time points and identified objects that had a length and width exceeding minimal values for a cell. The areas of these objects were signed, with negative values if a cell-shaped object disappeared and positive values if an object appeared at a given location. If the sum of these areas was less than −100 pixels, roughly the minimum size of a cell, we concluded that cell(s) had in fact left the field of view. Taking this sum accounted for cases where cells simply moved instead of detaching. Next, we simulated what the total microcolony area would have been had the cells remained and continued growing. Define the time-varying specific growth rate *µ*(*t*) of the departing cells as

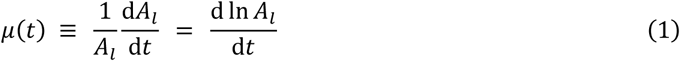

where *A_l_* is the area occupied by cells that left the field of view. Solve for *A_l_* gives

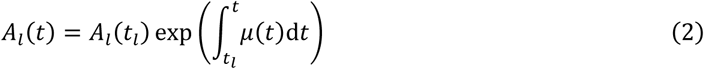

in which *t_l_* is the time when the cells left. We assumed that the departing cells would have grown at the same rate as the cells that remained. In other words,

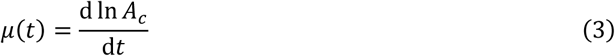

where *A_c_* is the measured area of the microcolony. We approximated this derivative using a first order forward difference. This derivative was inaccurate at the time of and immediately before any cell departure event. We replaced the value of the derivative at these times with the value closest in time that was unaffected by a departure. Then, accounting for all cell departures, the corrected microcolony area at time *t* was

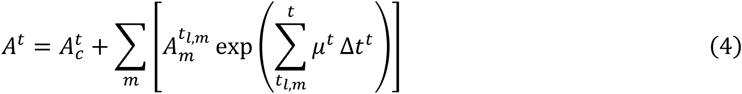

Here, the 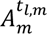 are the areas of all cells that departed at times *t*during the experiment. The measured microcolony areas and growth rates at time *t* are 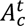 and *µ^t^* respectively. The time step Δ*t^t^* refers to time between image acquisitions at time *t*. Values of *A^t^* are plotted in Figure S8. For the -Ψ = 0 MPa treatment in panel A (nutrients present throughout, PEG never added), the corrected area increased exponentially as expected for microbial growth. Specific growth rates presented in the Results were computed by smoothing the corrected growth curves using locally weighted scatterplot smoothing and approximating the derivative as a first order forward difference. Smoothing was done separately for different experimental stages (i.e. adjustment, starvation, PEG stress, and recovery) to prevent smoothing over sudden changes in growth rate that occurred due to these transitions.

Any lysis event that results in a cell disappearing from view would also be detected using our approach. However, significant lysis was restricted to the -Ψ = 4 MPa treatment and occurred only after PEG was removed at the end of the experiments. Growth of the remaining cells was essentially zero so at worst, our estimates would miss cell death and indicate static biomass under these conditions. Growth estimation code is included in src/growth.py in Software S1.

### Modeling total fluorophore translation

To compare the extent of gene expression between treatments when PEG was present, we developed an integrative measure of reporter activity that accounted for several processes beyond fluorescent protein translation that affect fluorescence. Because mApple and mNeonGreen do not fluoresce until they have matured, we considered two pools of either protein: an invisible pool P and a fluorescent pool F. Light exposure also bleaches fluorescent proteins, causing their fluorescence to decrease over time. Finally, growth dilutes all proteins over a larger volume, decreasing their concentration. Assuming first order maturation and bleaching kinetics, the cytoplasmic concentrations of P and F vary according to these dynamics:

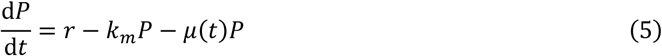

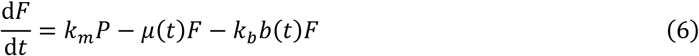

Here, *k_m_* is the maturation rate of protein P, *µ*(*t*) is the growth rate, and *k_b_* is the photobleaching rate of the matured, fluorescent protein F. The function *b*(*t*) is a sum of rectangular pulses with amplitude 1 and width equal to image exposure time beginning at each time when an image was taken. Finally, *r* is the translation rate, the quantity we aim to calculate, which depends on fluorescent protein gene transcription along with cellular translation capacity and other factors. We measured the cytoplasmic concentration of F in arbitrary fluorescence intensity units, so these units also apply to our estimated concentration of invisible protein P. We measured *k_b_* by placing exponential phase reporter cells in our PDMS well plates filled with SR media. Then, we took 200 images of the cells in rapid succession to induce photobleaching before new translation, protein maturation, or growth could occur. Finally, we estimated *k_b_* by fitting an exponential function to mApple and mNeonGreen fluorescence plotted against total excitation light exposure time (Figure S9). This value was 0.0180 ± 0.0001 (SE) s^−1^ for mApple and 0.0199 ± 0.0001 (SE) s^-1^ for mNeonGreen. Values of *k_m_* for mApple (1.4 hr^−1^) and mNeonGreen (4.2 hr^−1^) were taken from fpbase (69) and *µ*(*t*) was calculated as described above. For the pre-starved experiments, growth rates could only be calculated after PEG was removed and nutrients were restored so we assumed a growth rate of zero. We estimated the concentration of P at time *t* from *F* by discretizing equation 6 and solving for the *P_t_*:

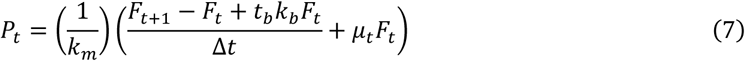

Here, *t_b_* is the exposure time, or duration of photobleaching, that occurs at each time point when an image is collected and Δ*t* is the time between images. Microcolony averaged values of *F* were smoothed via locally weighted scatterplot smoothing before they were substituted into this equation. The translation rate *r* was then computed by discretizing equation 5 and substituting the estimated values of *P_t_* into equation 5:

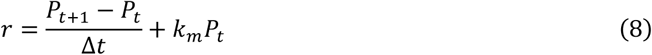

The total translation values for mApple and mNeonGreen that we report in Figure 6 were computed by integrating our estimates of *r* for each fluorophore over the time PEG was present. These values were slightly negative for some of the more stressful treatments, perhaps due to fluorophore leakage, degradation or other processes not included in our model. For this reason, we did not compute the ratio of integrated mNeonGreen to mApple fluorescence for the pre-starved cells, as both the numerator and denominator were near zero. This model is implemented in src/translation.py in Software S1.

**Figure 6.**
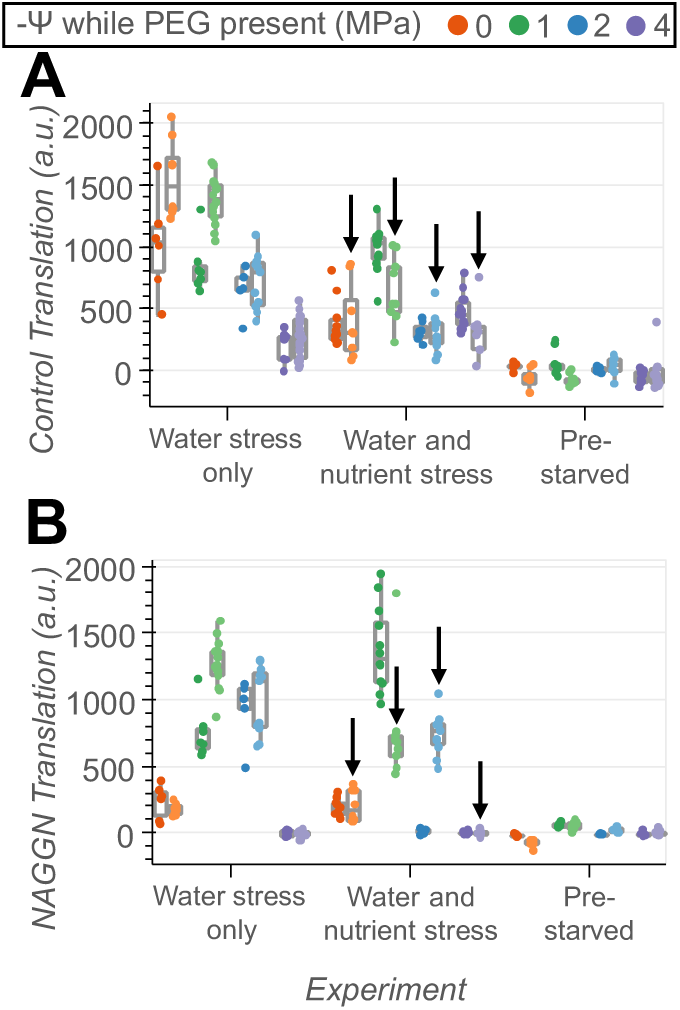
Comparing integrated fluorescent protein translation between nutrient and water stress treatments. (A) Total amount of mApple translation that occurred during PEG exposure across all experiments. Colors distinguish between water potentials. Lighter and darker shades represent different experimental replicates. For the experiments where water and nutrient stress began simultaneously (here labeled “Water and nutrient stress”), arrows point to the biological replicate that began with a slower growing inoculum. As these values were computed from fluorescence data (see Methods), they are expressed in arbitrary units. Points correspond to the microcolonies. (B) Total amount of mNeonGreen translation during the PEG treatments under control of the P_ggnA_ promoter.

## Supporting information

Figures S1-S9, Table S1, SI References

## Acknowledgments

We thank Reinaldo Alcalde for assistance with cloning and RNA sample preparation and Nate Glasser for assistance with computing RNA-seq fold changes. We also thank Alex Johnson for assistance with PDMS fabrication, Avi Flamholz for discussions on how to model translation from reporter fluorescence data, and all members of the Newman lab for their helpful input. We thank Saumya Saurabh for sharing the mNeonGreen plasmid. Finally, we thank Alessio Fragasso and Christine Jacobs-Wagner for sharing and demonstrating the PDMS well plate design. We prepared the PDMS well plates at the Resnick Ecology and Biosphere Engineering Facility, funded by the Resnick Sustainability Institute (RSI) at the California Institute of Technology (Caltech). This research was supported by Caltech’s Center for Environmental Microbial Interactions (Caldwell Fellowship awarded to JVK), a Caltech Division of Geological and Planetary Sciences Discovery Research Grant, and a 2022 Impact Grant from the RSI. JVK is a National Science Foundation Graduate Research Fellow (fellow ID 2021322073). GRS is a National Mah Jongg League Fellow of the Damon Runyon Cancer Research Foundation (DRG 2439-21).

## Notes

https://github.com/jkwiecinski0/NAGGNImageAnalysis

https://www.ncbi.nlm.nih.gov/bioproject/PRJNA1397269

### Competing Interest Statement

The authors have declared no competing interest.

## References

1. J. Huang, H. Yu, X. Guan, G. Wang, R. Guo, Accelerated dryland expansion under climate change. Nat. Clim. Change 6, 166–171 (2016).

2. B. R. Glick, Plant Growth-Promoting Bacteria: Mechanisms and Applications. Scientifica 2012, e963401 (2012).

3. J. I. Vílchez, C. García-Fontana, D. Román-Naranjo, J. González-López, M. Manzanera, Plant Drought Tolerance Enhancement by Trehalose Production of Desiccation-Tolerant Microorganisms. Front. Microbiol. 7 (2016).

4. J. P. Schimel, Life in Dry Soils: Effects of Drought on Soil Microbial Communities and Processes. Annu. Rev. Ecol. Evol. Syst. 49, 409–432 (2018).

5. H. F. Birch, The effect of soil drying on humus decomposition and nitrogen availability. Plant Soil 10, 9–31 (1958).

6. J. M. Wood, Osmosensing by Bacteria: Signals and Membrane-Based Sensors. Microbiol. Mol. Biol. Rev. 63, 230 (1999).

7. A. Dechesne, G. Wang, G. Gülez, D. Or, B. F. Smets, Hydration-controlled bacterial motility and dispersal on surfaces. Proc. Natl. Acad. Sci. 107, 14369–14372 (2010).

8. D. Or, B. F. Smets, J. M. Wraith, A. Dechesne, S. P. Friedman, Physical constraints affecting bacterial habitats and activity in unsaturated porous media – a review. Adv. Water Resour. 30, 1505–1527 (2007).

9. G. Wang, D. Or, A Hydration-Based Biophysical Index for the Onset of Soil Microbial Coexistence. Sci. Rep. 2, 881 (2012).

10. M. Potts, Desiccation tolerance of prokaryotes. Microbiol. Rev. 58, 755–805 (1994).

11. A. A. Malik, N. J. Bouskill, Drought impacts on microbial trait distribution and feedback to soil carbon cycling. Funct. Ecol. 36, 1442–1456 (2022).

12. R. D. Sleator, C. Hill, Bacterial osmoadaptation: the role of osmolytes in bacterial stress and virulence. FEMS Microbiol. Rev. 26, 49–71 (2002).

13. S. Manzoni, S. M. Schaeffer, G. Katul, A. Porporato, J. P. Schimel, A theoretical analysis of microbial eco-physiological and diffusion limitations to carbon cycling in drying soils. Soil Biol. Biochem. 73, 69–83 (2014).

14. D. S. Cayley, H. J. Guttman, M. T. Record, Biophysical characterization of changes in amounts and activity of Escherichia coli cell and compartment water and turgor pressure in response to osmotic stress. Biophys. J. 78, 1748–1764 (2000).

15. C. M. Boot, S. M. Schaeffer, J. P. Schimel, Static osmolyte concentrations in microbial biomass during seasonal drought in a California grassland. Soil Biol. Biochem. 57, 356–361 (2013).

16. P. H. Lebre, P. De Maayer, D. A. Cowan, Xerotolerant bacteria: surviving through a dry spell. Nat. Rev. Microbiol. 15, 285–296 (2017).

17. D.-E. Chang, D. J. Smalley, T. Conway, Gene expression profiling of Escherichia coli growth transitions: an expanded stringent response model. Mol. Microbiol. 45, 289–306 (2002).

18. P. Pletnev, I. Osterman, P. Sergiev, A. Bogdanov, O. Dontsova, Survival guide: Escherichia coli in the stationary phase. Acta Naturae 7, 22–33 (2015).

19. J. M. Navarro Llorens, A. Tormo, E. Martínez-García, Stationary phase in gram-negative bacteria. FEMS Microbiol. Rev. 34, 476–495 (2010).

20. B. H. J. Yap, S. A. Crawford, R. R. Dagastine, P. J. Scales, G. J. O. Martin, Nitrogen deprivation of microalgae: effect on cell size, cell wall thickness, cell strength, and resistance to mechanical disruption. J. Ind. Microbiol. Biotechnol. 43, 1671–1680 (2016).

21. W. Liang, et al., Adaptation to Long-Term Nitrogen Starvation in a Biocrust-Derived Microalga Vischeria sp. WL1: Insights into Cell Wall Features and Desiccation Resistance. Microorganisms 13, 903 (2025).

22. D. E. Jenkins, S. A. Chaisson, A. Matin, Starvation-induced cross protection against osmotic challenge in Escherichia coli. J. Bacteriol. 172, 2779–2781 (1990).

23. D. Tashyreva, J. Elster, Effect of nitrogen starvation on desiccation tolerance of Arctic Microcoleus strains (cyanobacteria). Front. Microbiol. 6 (2015).

24. C. C. Cleveland,. Liptzin, C:N:P stoichiometry in soil: is there a “Redfield ratio” for the microbial biomass? Biogeochemistry 85, 235–252 (2007).

25. L. J. Halverson, M. K. Firestone, Differential Effects of Permeating and Nonpermeating Solutes on the Fatty Acid Composition of Pseudomonas putida. Appl. Environ. Microbiol. 66, 2414–2421 (2000).

26. M. Geisler, et al., Analysis of high-molecular weight polyethylene glycol degradation by *Pseudomonas* sp. Polym. Degrad. Stab. 232, 111144 (2025).

27. L. T. Smith, G. M. Smith, An osmoregulated dipeptide in stressed Rhizobium meliloti. J. Bacteriol. 171, 4714–4717 (1989).

28. B. Sagot, et al., Osmotically induced synthesis of the dipeptide N-acetylglutaminylglutamine amide is mediated by a new pathway conserved among bacteria. Proc. Natl. Acad. Sci. 107, 12652–12657 (2010).

29. M. Kurz, A. Y. Burch, B. Seip, S. E. Lindow, H. Gross, Genome-Driven Investigation of Compatible Solute Biosynthesis Pathways of Pseudomonas syringae pv. syringae and Their Contribution to Water Stress Tolerance. Appl. Environ. Microbiol. 76, 5452–5462 (2010).

30. B. C. Freeman, et al., Physiological and Transcriptional Responses to Osmotic Stress of Two Pseudomonas syringae Strains That Differ in Epiphytic Fitness and Osmotolerance. J. Bacteriol. 195, 4742–4752 (2013).

31. W.-S. Chang, et al., Alginate Production by Pseudomonas putida Creates a Hydrated Microenvironment and Contributes to Biofilm Architecture and Stress Tolerance under Water-Limiting Conditions. J. Bacteriol. 189, 8290–8299 (2007).

32. J. McWilliams, Pseudomonas/Brachypodium as a Model System for Studying Rhizosphere Plant Microbe Interactions Under Water Stress. Masters Theses (2018).

33. A. Fragasso, T. Schlechtweg, W.-H. Lin, A. E. Barron, C. Jacobs-Wagner, Time-resolved phenotyping at subcellular resolution reveals shared principles and key trade-offs across antimicrobial peptide activities. [Preprint] (2025). Available at: https://www.biorxiv.org/content/10.1101/2025.04.10.648262v1 [Accessed 29 October 2025].

34. S. Vadia, P. A. Levin, Growth rate and cell size: a re-examination of the growth law. Curr. Opin. Microbiol. 24, 96–103 (2015).

35. D. Metze, et al., Microbial growth under drought is confined to distinct taxa and modified by potential future climate conditions. Nat. Commun. 14, 5895 (2023).

36. J. T. Lennon, B. K. Lehmkuhl, A trait-based approach to bacterial biofilms in soil. Environ. Microbiol. 18, 2732–2742 (2016).

37. D. Or, S. Phutane, A. Dechesne, Extracellular Polymeric Substances Affecting Pore-Scale Hydrologic Conditions for Bacterial Activity in Unsaturated Soils. Vadose Zone J. 6, 298–305 (2007).

38. S. J. Blazewicz, et al., Taxon-specific microbial growth and mortality patterns reveal distinct temporal population responses to rewetting in a California grassland soil. ISME J. 14, 1520–1532 (2020).

39. S. Evans, U. Dieckmann, O. Franklin, C. Kaiser, Synergistic effects of diffusion and microbial physiology reproduce the Birch effect in a micro-scale model. Soil Biol. Biochem. 93, 28–37 (2016).

40. A. Canarini, et al., Ecological memory of recurrent drought modifies soil processes via changes in soil microbial community. Nat. Commun. 12, 5308 (2021).

41. C. R. Warren, Do microbial osmolytes or extracellular depolymerisation products accumulate as soil dries? Soil Biol. Biochem. 98, 54–63 (2016).

42. E. W. Slessarev, et al., Cellular and extracellular C contributions to respiration after wetting dry soil. Biogeochemistry 147, 307–324 (2020).

43. S. Jamal, et al., Relationship between functional activity and protein stability in the presence of all classes of stabilizing osmolytes. FEBS J. 276, 6024–6032 (2009).

44. H. Tapia, D. E. Koshland, Trehalose Is a Versatile and Long-Lived Chaperone for Desiccation Tolerance. Curr. Biol. 24, 2758–2766 (2014).

45. H. Tapia, L. Young, D. Fox, C. R. Bertozzi, D. Koshland, Increasing intracellular trehalose is sufficient to confer desiccation tolerance to Saccharomyces cerevisiae. Proc. Natl. Acad. Sci. 112, 6122–6127 (2015).

46. R. Hengge-Aronis, W. Klein, R. Lange, M. Rimmele, W. Boos, Trehalose synthesis genes are controlled by the putative sigma factor encoded by rpoS and are involved in stationary-phase thermotolerance in Escherichia coli. J. Bacteriol. 173, 7918–7924 (1991).

47. B. R. Parry, et al., The Bacterial Cytoplasm Has Glass-like Properties and Is Fluidized by Metabolic Activity. Cell 156, 183–194 (2014).

48. M. Potts, Desiccation tolerance: a simple process? Trends Microbiol. 9, 553–559 (2001).

49. H. Shi, et al., Starvation induces shrinkage of the bacterial cytoplasm. Proc. Natl. Acad. Sci. 118, e2104686118 (2021).

50. D. M. Weller, R. J. Cook, Suppression of Take-All of Wheat by Seed Treatments with Fluorescent Pseudomonads. Phytopathology® 115, 463–469 (1983).

51. L. S. Thomashow, D. M. Weller, Role of a phenazine antibiotic from Pseudomonas fluorescens in biological control of Gaeumannomyces graminis var. tritici. J. Bacteriol. 170, 3499–3508 (1988).

52. O. V. Mavrodi, D. V. Mavrodi, J. A. Parejko, L. S. Thomashow, D. M. Weller, Irrigation Differentially Impacts Populations of Indigenous Antibiotic-Producing Pseudomonas spp. in the Rhizosphere of Wheat. Appl. Environ. Microbiol. 78, 3214–3220 (2012).

53. B. L. Mortensen, S. Rathi, W. J. Chazin, E. P. Skaar, Acinetobacter baumannii Response to Host-Mediated Zinc Limitation Requires the Transcriptional Regulator Zur. J. Bacteriol. 196, 2616–2626 (2014).

54. B. Langmead, S. L. Salzberg, Fast gapped-read alignment with Bowtie 2. Nat. Methods 9, 357–359 (2012).

55. Y. Liao, G. K. Smyth, W. Shi, featureCounts: an efficient general purpose program for assigning sequence reads to genomic features. Bioinformatics 30, 923–930 (2014).

56. M. I. Love, W. Huber, S. Anders, Moderated estimation of fold change and dispersion for RNA-seq data with DESeq2. Genome Biol. 15, 550 (2014).

57. M. B. Elowitz, S. Leibler, A synthetic oscillatory network of transcriptional regulators. Nature 403, 335–338 (2000).

58. N. C. Shaner, et al., A bright monomeric green fluorescent protein derived from Branchiostoma lanceolatum. Nat. Methods 10, 407–409 (2013).

59. W. G. Miller, J. H. Leveau, S. E. Lindow, Improved gfp and inaZ broad-host-range promoter-probe vectors. Mol. Plant-Microbe Interact. MPMI 13, 1243–1250 (2000).

60. K.-H. Choi, A. Kumar, H. P. Schweizer, A 10-min method for preparation of highly electrocompetent *Pseudomonas aeruginosa* cells: Application for DNA fragment transfer between chromosomes and plasmid transformation. J. Microbiol. Methods 64, 391–397 (2006).

61. X. Dai, et al., Slowdown of Translational Elongation in Escherichia coli under Hyperosmotic Stress. mBio 9, e02375–17 (2018).

62. N. C. Shaner, et al., Improving the photostability of bright monomeric orange and red fluorescent proteins. Nat. Methods 5, 545–551 (2008).

63. J. Meisner, J. B. Goldberg, The Escherichia coli rhaSR-PrhaBAD Inducible Promoter System Allows Tightly Controlled Gene Expression over a Wide Range in Pseudomonas aeruginosa. Appl. Environ. Microbiol. 82, 6715–6727 (2016).

64. F. M. Ausubel, Current Protocols in Molecular Biology (Greene Pub. Associates and Wiley-Interscience, 1987).

65. K.-H. Choi, et al., A Tn7-based broad-range bacterial cloning and expression system. Nat. Methods 2, 443–448 (2005).

66. L. Lambertsen, C. Sternberg, S. Molin, Mini-Tn7 transposons for site-specific tagging of bacteria with fluorescent proteins. Environ. Microbiol. 6, 726–732 (2004).

67. T. W. Lo, K. J. Cutler, H. J. Choi, P. A. Wiggins, OmniSegger: A time-lapse image analysis pipeline for bacterial cells. PLOS Comput. Biol. 21, e1013088 (2025).

68. K. J. Cutler, et al., Omnipose: a high-precision morphology-independent solution for bacterial cell segmentation. Nat. Methods 19, 1438–1448 (2022).

69. T. J. Lambert, FPbase: a community-editable fluorescent protein database. Nat. Methods 16, 277–278 (2019).

70. National Ecological Observatory Network (NEON), Soil water content and water salinity (DP1.00094.001). National Ecological Observatory Network (NEON). 10.48443/qhmt-hh62. Deposited 2025.

71. R. J. Millington, J. P. Quirk, Permeability of porous solids. Trans. Faraday Soc. 57, 1200–1207 (1961).

72. M. Th. van Genuchten, A Closed-form Equation for Predicting the Hydraulic Conductivity of Unsaturated Soils. Soil Sci. Soc. Am. J. 44, 892–898 (1980).

73. National Ecological Observatory Network (NEON), Soil physical and chemical properties, Megapit (DP1.00096.001). National Ecological Observatory Network (NEON). 10.48443/0hnd-yj57. Deposited 2025.

74. R. F. Carsel, R. S. Parrish, Developing joint probability distributions of soil water retention characteristics. Water Resour. Res. 24, 755–769 (1988).

