## Supplementary material for "Starvation enhances bacterial survival through drying and rewetting at the single cell level": Figures S1-S9, Table S1, SI References

**This PDF file includes:**

Figures S1 to S9
Table S1
SI References

**Other supporting materials for this manuscript include the following:**

Table S2
Datasets S1 to S3
Software S1

**Figures**

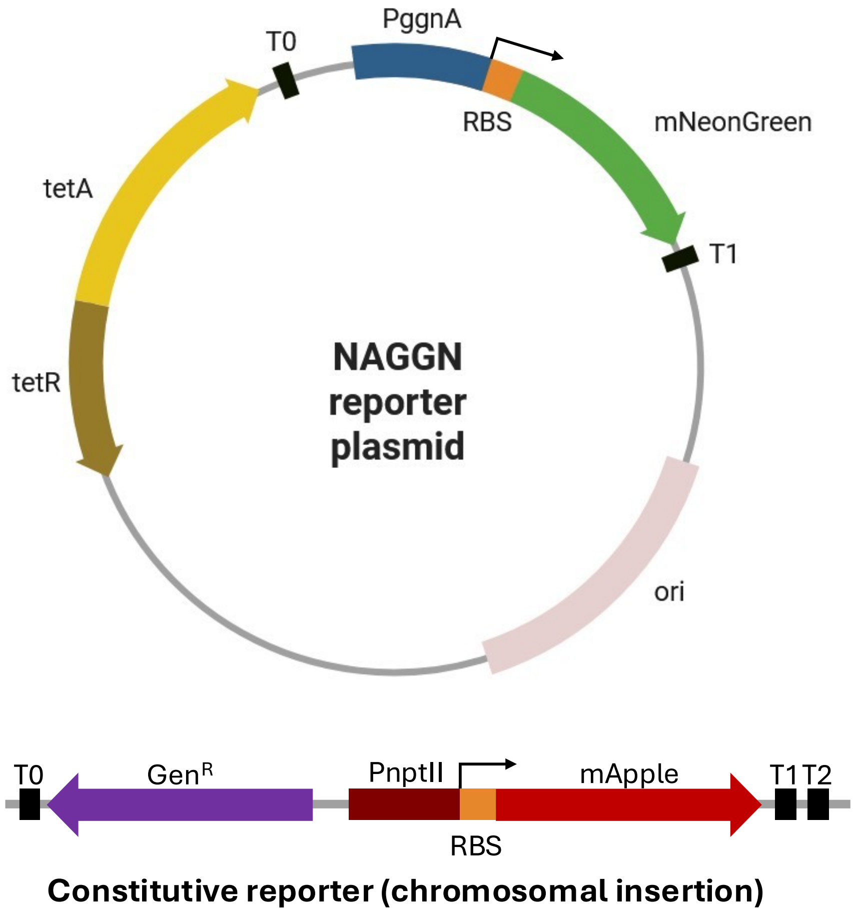


**Fig. S1.** Design of transcriptional reporter used to monitor NAGGN biosynthesis and overall gene expression. The gene for the fluorescent protein mNeonGreen under control of the $P_{ggnA}$ promoter was cloned into the pPROBETT plasmid backbone, a vector that contains an origin of replication recognized by pseudomonads and with a tetracycline resistance marker. T0 and T1 are transcriptional terminators, and the RBS is a synthetic ribosome binding site. Black, right-angled arrows show transcription start sites. The constitutive promoter $P_{\mathbf{nptII}}$ was added to the chromosome at the *att*Tn7 site and drove expression of the gene for the fluorescent protein mApple. The gentamycin resistance cassette included in this construct was used for selection while cloning.

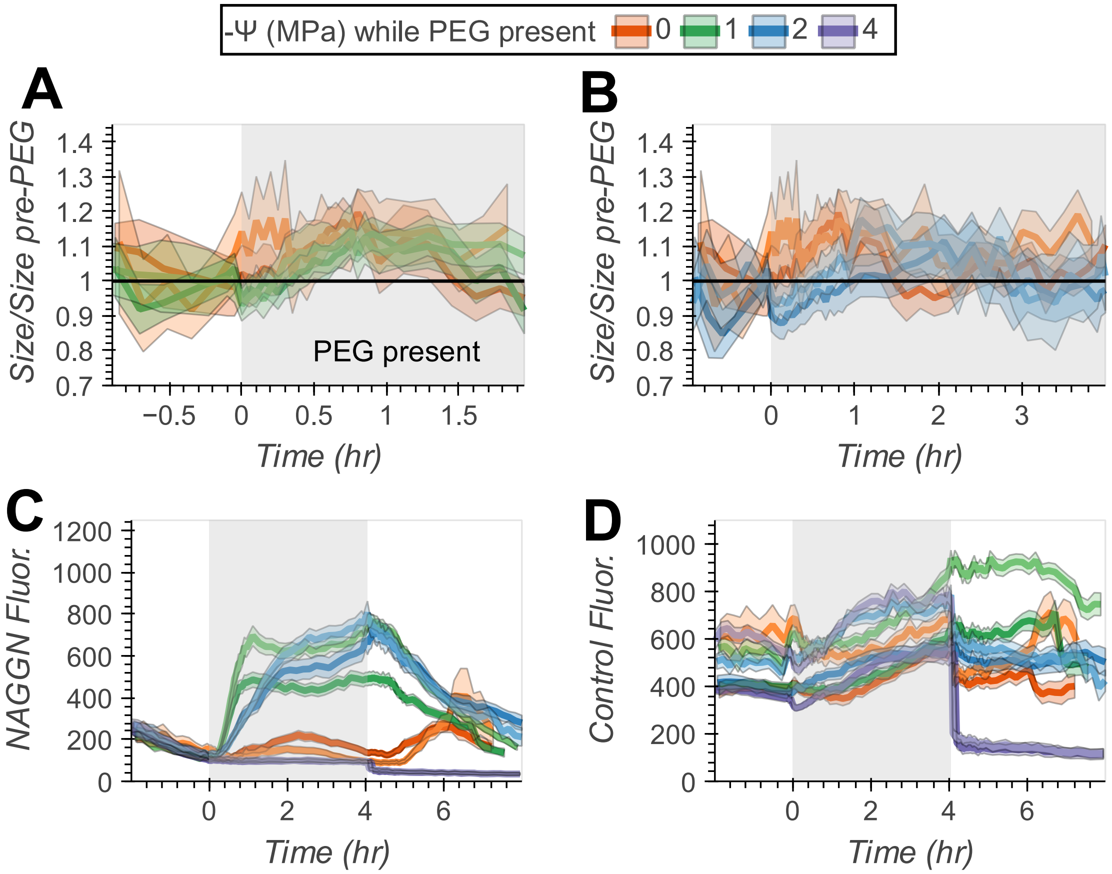


**Fig. S2.** Supplementary data for the experiments with water stress only. (A) The normalized size of reporter cells from the -Ψ = 0 MPa and 1 MPa treatments. Lines show the mean cell size at each time point normalized to the mean size in the 30 minutes leading up to PEG addition. Colors show different water potentials, and the darker and lighter shades distinguish between replicate experiments. Ranges show 95% confidence intervals. During PEG exposure (grey shading), there was an average of 44-94 cells present for the water potentials shown, with numbers varying between water potentials and replicates. Complete cell counts for the entire time series are given in Figure S6. (B) Normalized size of reporter cells in the -Ψ = 0 and 2 MPa treatments. During PEG exposure, there was an average of 33-48 cells present for the water potentials shown. (C) Time series of mNeonGreen fluorescence under control of the $P_{ggnA}$ promoter, a proxy for NAGGN biosynthesis. Lines show mean fluorescence across all segmentation masks, including those with multiple cells. There were on average 46-204 segmentation masks present during the PEG treatments, and a complete record of the number of masks is given in Figure S5. (D) Constitutive mApple, or control, fluorescence under control of the $P_{nptII}$ promoter, with the same number of masks used to compute averages as in panel C.

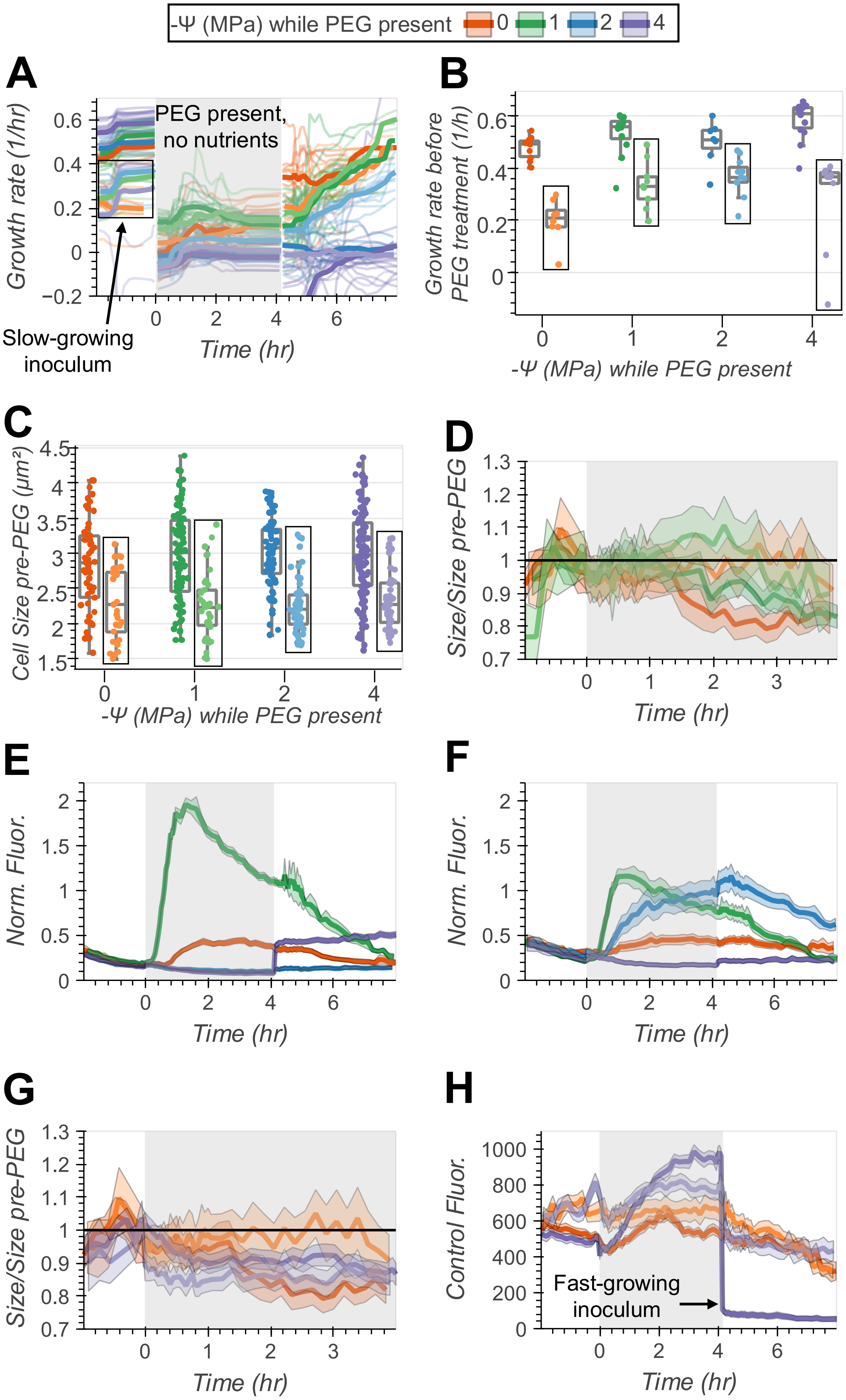
-Ψ (MPa) while PEG present 0 1 2 4
A
B
Growth rate (1/hr)
PEG present,
no nutrients
Slow-growing
inoculum
Time (hr)
Growth rate before
PEG treatment (1/h)
-Ψ (MPa) while PEG present
C
D
Cell Size pre-PEG (μm²)
-Ψ (MPa) while PEG present
Size/Size pre-PEG
Time (hr)
E
F
Norm. Fluor.
Time (hr)
G
H
Size/Size pre-PEG
Time (hr)
Control Fluor.
Fast-growing
inoculum

**Fig. S3.** Supplementary data for experiments where nutrients and water were simultaneously removed. (A) Time series of growth rates. In one of the two biological replicates (lighter colors), the growth rate was lower before PEG was added and nutrients were removed (grey shading). Bold lines show the mean growth rate across all microcolonies and faint lines show growth rates for individual microcolonies. The number of microcolonies measured at each time point is displayed in Figure S7. (B) Growth rate prior to PEG addition and nutrient withdrawal. Points show the mean growth rate for each microcolony in the hour leading up to PEG addition. Colors show the water potentials that were imposed once PEG was added. Boxes indicate microcolonies that originated from the slower growing inoculum. (C) Comparison of absolute cell sizes in the slow and fast-growing replicates at the end of the adjustment period immediately before PEG was added and nutrients were removed. (D) The normalized size of reporter cells from the $-\Psi = 0$ MPa and 1 MPa treatments. Lines show the mean cell size at each time point normalized to the mean size in the 30 minutes leading up to PEG addition. Colors show different water potentials, and the darker and lighter shades distinguish between replicate experiments. Ranges show 95% confidence intervals. During PEG exposure and nutrient deprivation (grey shading), there was an average of 27-125 cells present for the water potentials shown, with numbers varying between water potentials and replicates. Complete cell counts for the entire time series are given in Figure S6. (E) NAGGN reporter mNeonGreen fluorescence normalized by constitutive mApple fluorescence from the experimental replicate that began with a faster growing inoculum. Lines represent the mean normalized fluorescence across all segmentation masks, including those with multiple cells. During PEG exposure, there was an average of 107-253 masks present per water potential and replicate. Complete mask counts for the entire time series are in Figure S5. (F) Normalized fluorescence from the replicate with a slower growing inoculum. During PEG exposure, there was an average of 45-72 masks present per water potential and replicate. (G) Normalized size of cells from the $-\Psi = 0$ and 4 MPa treatments in both replicates. During PEG exposure, there was an average of 34-229 cells present for the water potentials shown. (H) Raw constitutive mApple fluorescence from the $-\Psi = 0$ and 4 MPa treatments averaged across all segmentation masks. During PEG exposure, there was an average of 45-253 masks present for the water potentials shown. Note the sharp decrease in fluorescence once PEG was removed and nutrients were restored for cells that originated from the fast-growing inoculum.

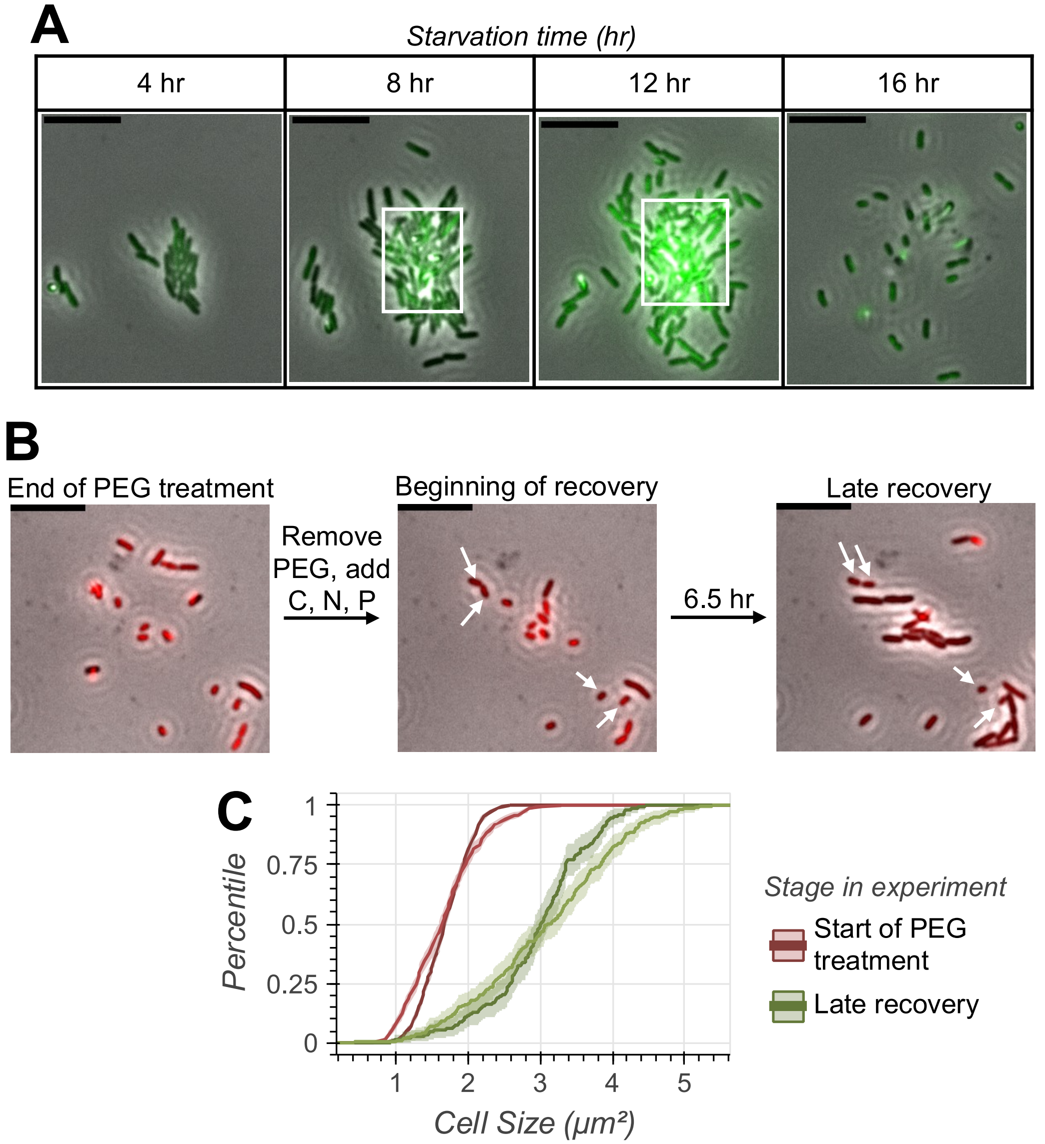


**Fig. S4.** Supplementary data for the pre-starved experiments. (A) Images of reporter cells during starvation. Times in the top column are the number of hours that cells were exposed to the starvation medium. The microcolony shown more than doubled in size between 4 and 8 hours of starvation but enlarged by much less between 8 and 12 hours. Cells also began growing on top of each other after ~7 hours of starvation, especially within the white rectangles superimposed on the 8 and 12 hour images. Finally, most cells detached from the Petri dish between 12 and 16 hours. Green represents the mNeonGreen fluorescence intensity, and all images are on the same intensity scale. Scale bars are 10 μm. (B) Images of reporter cells at the end of the PEG treatment, immediately after PEG was removed and nutrients were restored, and 6.5 hours into the recovery period. The red overlay is mApple fluorescence, showing that cells remained intact throughout these transitions. White arrows in the images during recovery show cells that did not resume growth. Scale bars are 10μm and all images are on the same red fluorescence intensity scale. (C) Distributions of cell sizes from the -Ψ = 4 MPa treatment at different times. The red distributions show sizes 30 minutes after PEG addition. The green distributions show the size at $t$ = 10.5 hr, 6.5 hours after PEG was removed and nutrients were restored. This matches the time that the last image in S4B was taken. Light and dark shades again distinguish between replicate experiments. Ranges show 95% confidence intervals computed via bootstrapping with 10000 bootstrap replicates.

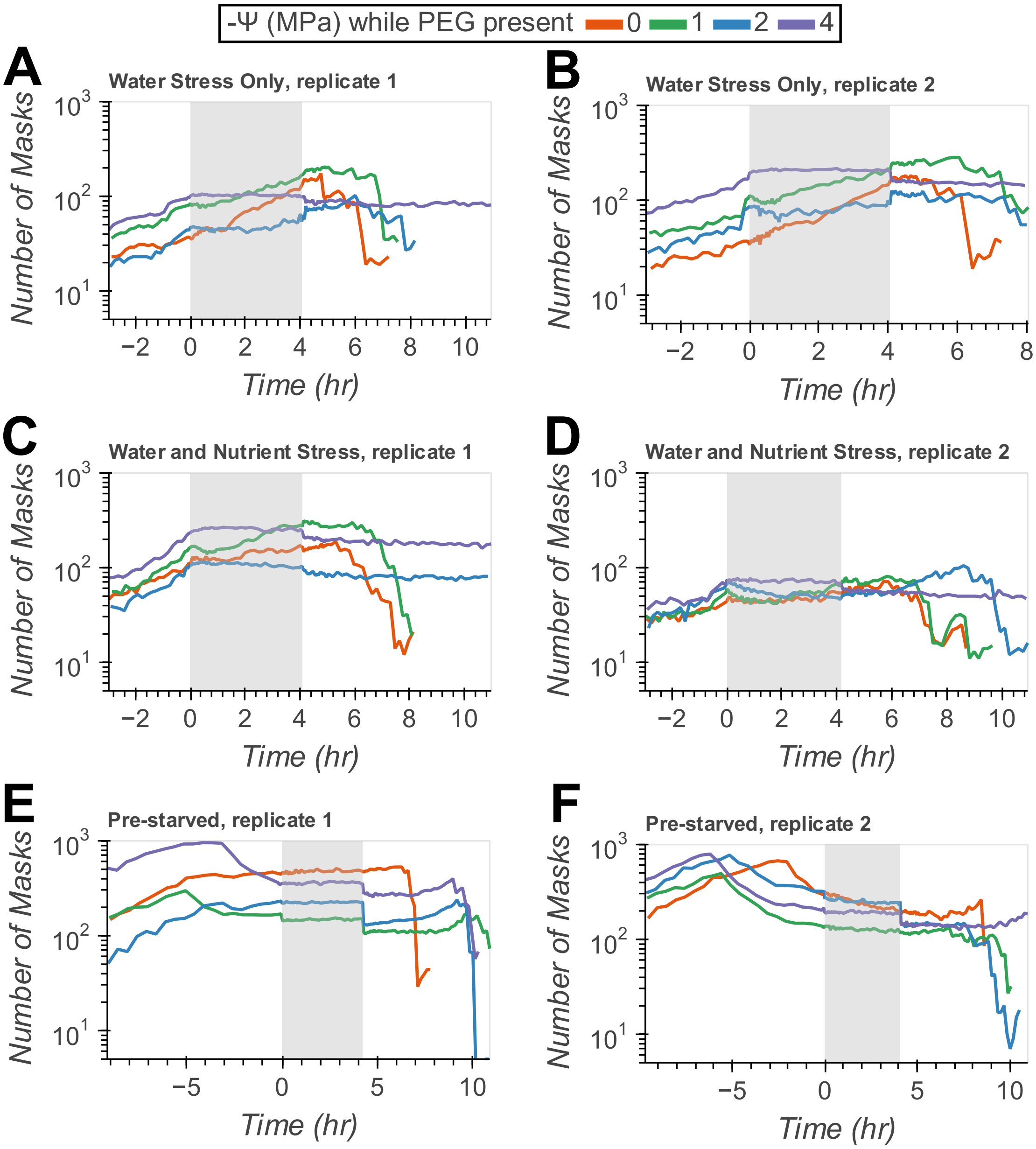


**Fig. S5.** Number of unfiltered segmentation masks used to compute fluorescence means. Grey shading shows when PEG was present. For the water and nutrient stress experiments (C-D), cells in replicate 1 grew faster than cells in replicate 2 prior to PEG addition.

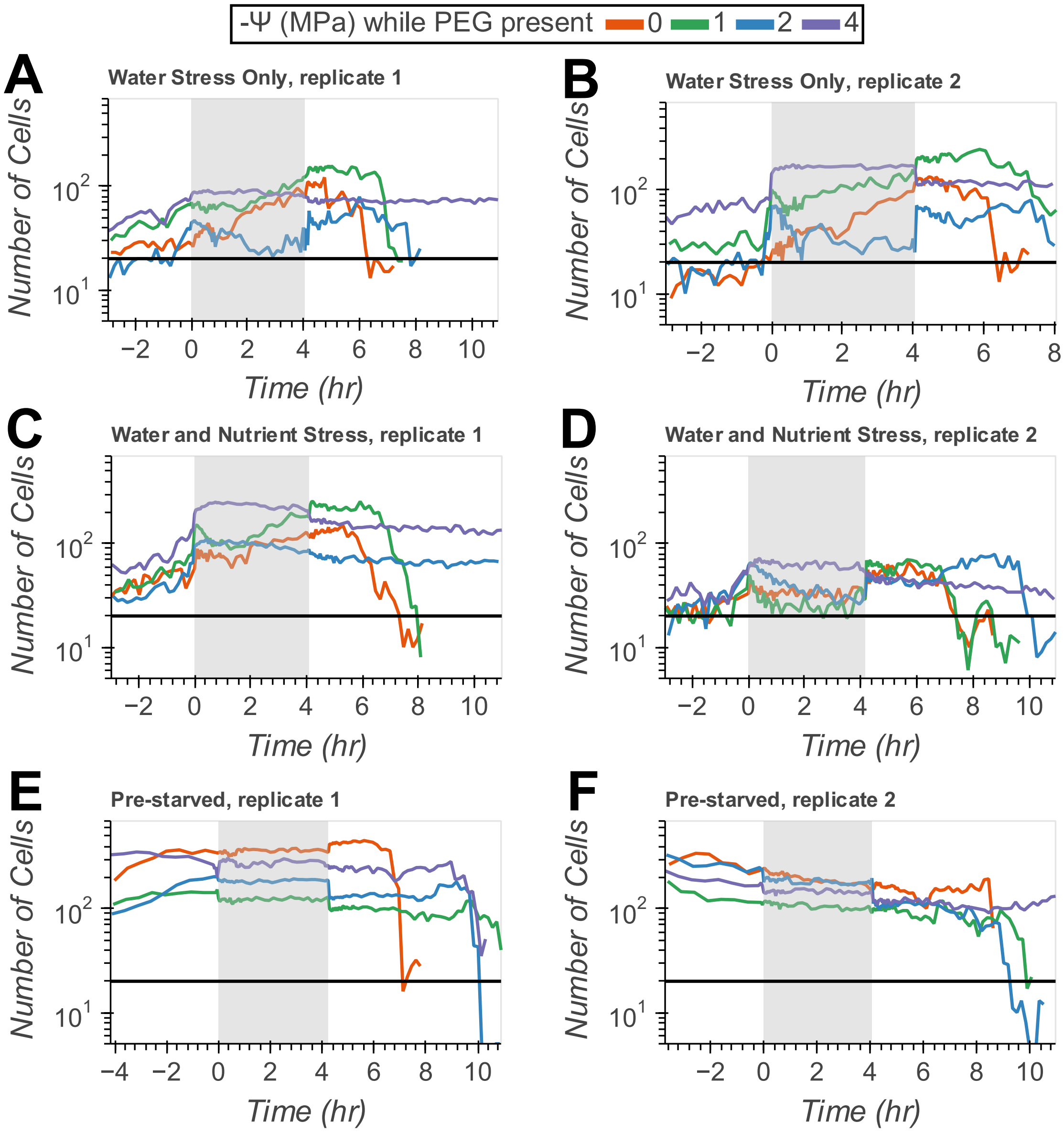


**Fig. S6.** Number of filtered segmentation masks containing one cell that were used to compute mean cell sizes. Grey shading shows when PEG was present. A line is drawn at n=20 to indicate that there were at least this many masks with only one cell present throughout the PEG treatments in all conditions. For the water and nutrient stress experiments (C-D), cells in replicate 1 grew faster than cells in replicate 2 prior to PEG addition.

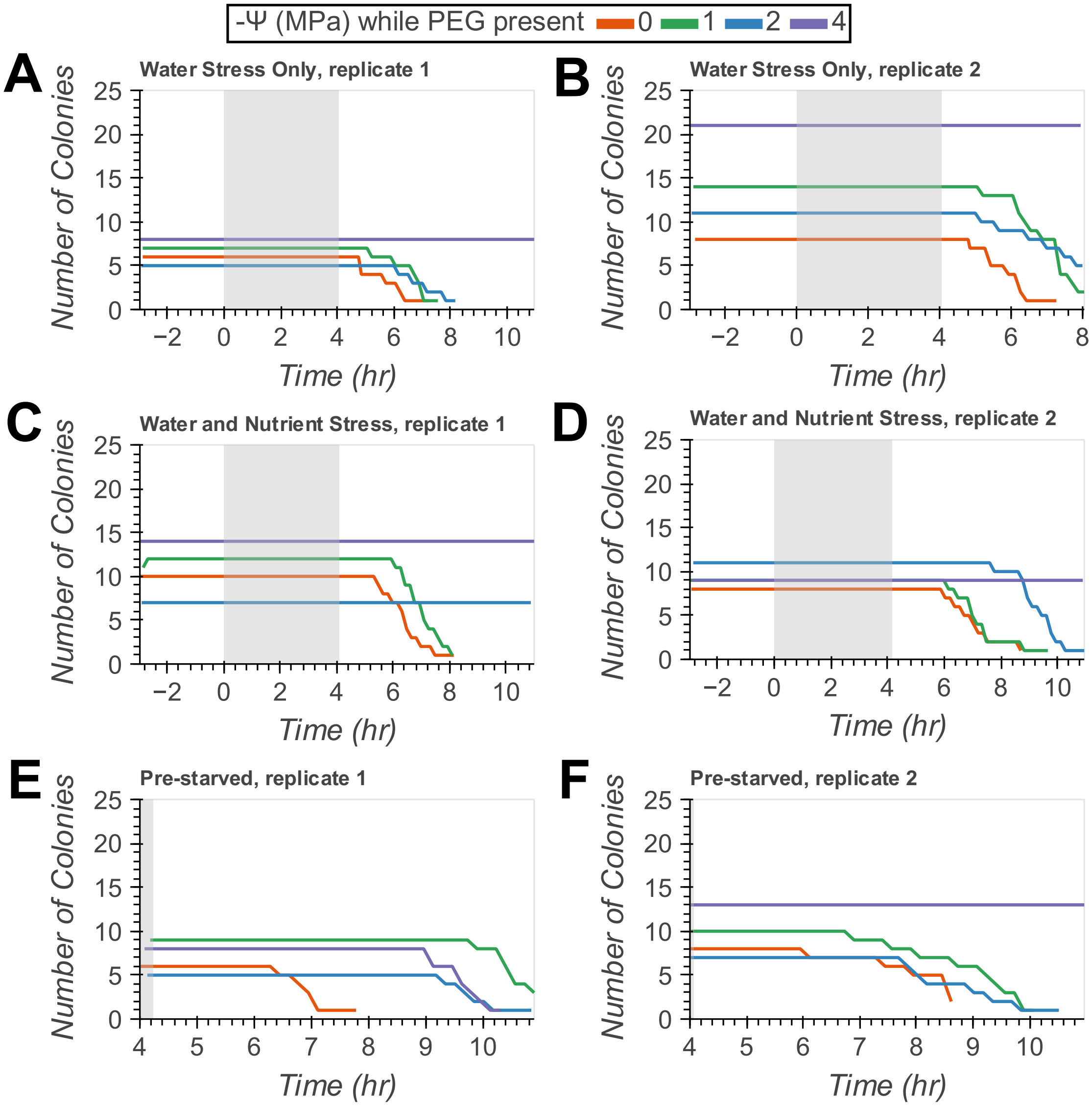


**Fig. S7.** Number of microcolonies for which we estimated growth and translation rates. Grey shading shows when PEG was preset. Counts are shown over the time range for which the growth rate could be calculated. For the water and nutrient stress experiments (C-D), cells in replicate 1 grew faster than cells in replicate 2 prior to PEG addition.

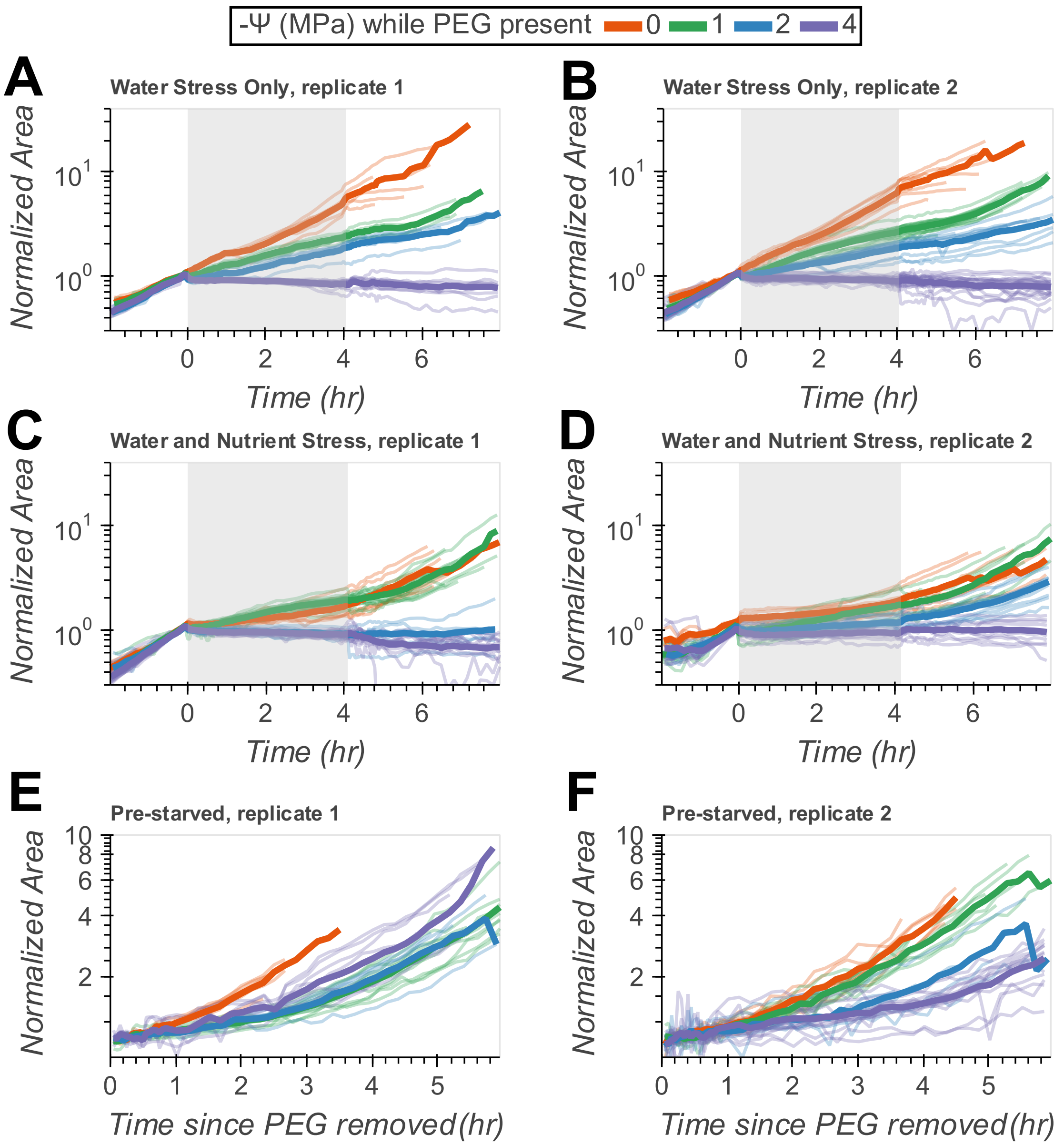


**Fig. S8.** Growth curves. (A, B) Corrected microcolony areas for the experiments with water stress only. Bold lines show the mean area across all microcolonies and faint lines show the area for individual microcolonies. Lines are colored by water potential. Areas are normalized to the mean area in the 30 minutes prior to PEG addition. Gray shading shows when PEG was present. (C, D) Corrected microcolony areas for the experiments in which nutrients and water were removed simultaneously at *t* = 0 hr. Cells in replicate 1 grew faster than cells in replicate 2 prior to PEG addition. (E, F) Growth of pre-starved microcolonies once PEG was removed and nutrients were restored. Areas are normalized to the mean area in the first 30 minutes after PEG was removed.

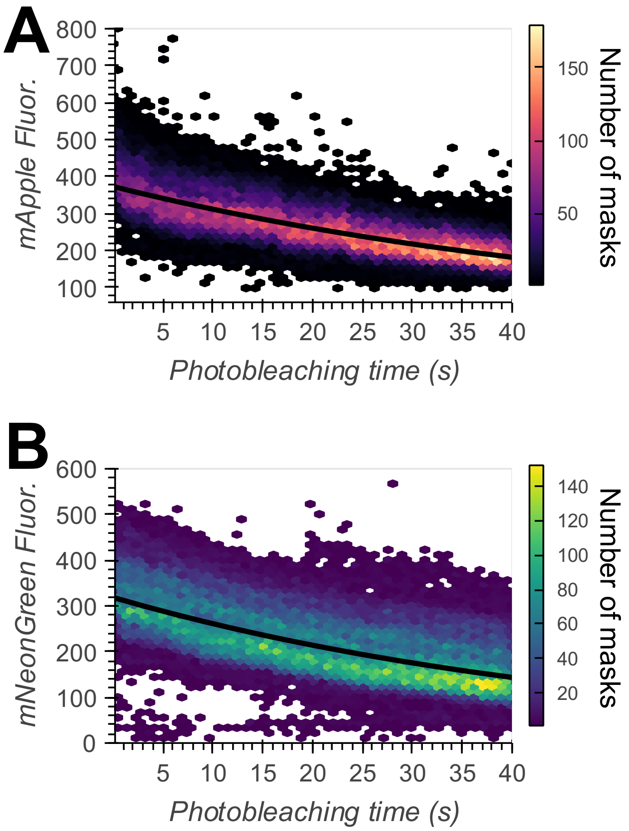


**Fig. S9.** Photobleaching time series. (A) mApple fluorescence vs. photobleaching time. Color shading shows the number of segmentation masks falling within each bin. Masks contained one or more cells. The black line shows the exponential function fit to the data to derive the photobleaching rate constant. (B) mNeonGreen photobleaching time series.

### Tables

**Table S1. Media Compositions**

| SR Medium | |
|---|---|
| **Component** | **Concentration (mM)** |
| $NH_4Cl$ | 8.57 |
| $K_2HPO_4$ | 0.29 |
| $KH_2PO_4$ | 0.71 |
| $MgSO_4$ | 0.41 |
| $CaCl_2$ | 0.68 |
| Glucose | 10.00 |
| Casamino acids | 0.05% v/v |
| MOPS | 71.50 |
| NaMOPS | 28.50 |
| KCl | 10.00 |
| Aquil Trace Metal Mix (1) | 1X strength |

| SRM Medium | |
|---|---|
| **Component** | **Concentration (mM)** |
| $MgSO_4$ | 0.41 |
| $CaCl_2$ | 0.68 |
| MOPS | 80.65 |
| NaMOPS | 32.14 |
| KCl | 11.29 |
| NaCl | 5.21 |
| Aquil Trace Metal Mix (1) | 1X strength |

Composition of SR and SRM media. For both media, the total osmolarity excluding any contribution from casamino acids or the trace metal mix is 180.79 mM and the ionic strength is 52.32 mM. Phosphate salts and MOPS buffer were added in proportions to set the pH at 6.8.

**Table S2 (separate file).** Table of strains, plasmids, and primers. In primer sequences, capital letters show annealing regions and lowercase letters show homologous regions for Gibson assembly. References for plasmid sources are listed in the SI References.

**Dataset S1 (separate file).** Fold changes and p-values for all genes in the RNA-seq experiment. In the spreadsheet, Locus is the locus tags for genes in *P. synxantha*'s genome. baseMeans is the average number of reads mapped to a locus across all samples. lfcse is the standard error in the log2FoldChange and stat contains values of the Wald test statistic used to compute p values. padj is the p value adjusted using the Benjamini and Hochberg method. These columns were generated by DESeq2. We also provide GenBank annotations where available for all locus tags.

**Dataset S2.** Raw reads from RNA-seq experiment available on the NCBI Sequence Read Archive under accession number PRJNA1397269.

**Dataset S3 (separate file).** Annotated sequence of reporter plasmid.

**Software S1.** Code used to perform image analysis and generate all figures available at https://github.com/jkwiecinski0/NAGGNImageAnalysis.

**SI References**


1. D. E. Martocello, F. M. M. Morel, D. L. McRose, H-Aquil: a chemically defined cell culture medium for trace metal studies in Vibrios and other marine heterotrophic bacteria. *Biometals* **32**, 819–828 (2019).
2. J. McWilliams, Pseudomonas/Brachypodium as a Model System for Studying Rhizosphere Plant Microbe Interactions Under Water Stress. *Master's Theses* (2018).
3. R. Chawla, *et al.*, Reentrant DNA shells tune polyphosphate condensate size. *Nat Commun* **15**, 9258 (2024).
4. D. W. Basta, M. Bergkessel, D. K. Newman, Identification of Fitness Determinants during Energy-Limited Growth Arrest in Pseudomonas aeruginosa. *mBio* (2017). https://doi.org/10.1128/mbio.01170-17.
5. M. A. Jacobs, *et al.*, Comprehensive transposon mutant library of Pseudomonas aeruginosa. *Proceedings of the National Academy of Sciences* **100**, 14339–14344 (2003).
6. L. Lambertsen, C. Sternberg, S. Molin, Mini-Tn7 transposons for site-specific tagging of bacteria with fluorescent proteins. *Environmental Microbiology* **6**, 726–732 (2004).
